# Origin and evolutionary trajectories of brown algal sex chromosomes

**DOI:** 10.1101/2024.01.15.575685

**Authors:** Josué Barrera-Redondo, Agnieszka P. Lipinska, Pengfei Liu, Erica Dinatale, Guillaume Cossard, Kenny Bogaert, Masakazu Hoshino, Rory J. Craig, Komlan Avia, Goncalo Leiria, Elena Avdievich, Daniel Liesner, Rémy Luthringer, Olivier Godfroy, Svenja Heesch, Zofia Nehr, Loraine Brillet-Guéguen, Akira F. Peters, Galice Hoarau, Gareth Pearson, Jean-Marc Aury, Patrick Wincker, France Denoeud, J Mark Cock, Fabian B. Haas, Susana M Coelho

## Abstract

Sex chromosomes fall into three classes: XX/XY, ZW/ZZ and U/V systems. The rise, evolution and demise of U/V systems has remained an evolutionary enigma. Here, we analyse genomes spanning the entire brown algal phylogeny to decipher their sex-determination evolutionary history. U/V sex chromosomes emerged between 450 and 224 million years ago, when a region containing the pivotal male-determinant *MIN* located in a discrete region in proto-U and proto-V chromosomes ceased recombining. Over time, nested inversions led to step-wise expansions of the sex locus, accompanying increasing morphological complexity and sexual differentiation of brown seaweeds. Unlike XX/XY and ZW/ZZ, brown algal U/V evolve mainly by gene gain, showing minimal degeneration. They are structurally dynamic and act as genomic ‘cradles’ fostering the birth of new genes, potentially from ancestrally non coding sequences. Our analyses demonstrate that hermaphroditism arose from ancestral males that acquired U-specific genes by ectopic recombination, and that in the transition from a U/V to an XX/XY system, V-specific genes moved down the genetic hierarchy of sex determination. Both events lead to the demise of U and V and erosion of their specific genomic characteristics. Taken together, our findings offer a comprehensive model of U/V sex chromosome evolution.

## INTRODUCTION

Sexual reproduction, a feature in almost all eukaryotes, enables species to survive environmental challenges by increasing genetic variation^1^. While fundamental mechanisms of meiosis and fertilization are largely conserved, the paths instructing acquisition of male or female identities vary widely across species^2,3^. Sex chromosomes carry a sex-determining region (SDR)^4^ encoding genetic factors that instruct sex identity and which does not undergo recombination in the heterogametic sex (XY or ZW)^1^. Sex chromosomes have independently and repeatedly evolved from autosomes (*i.e.*, any chromosome that is not a sex chromosome) and are subjected to specific evolutionary forces, such as differential selection in male and females, asymmetrical sheltering of deleterious mutations, hemizygosity, meiotic silencing and dosage compensation^4^. They also play prominent roles in speciation and adaptation^1,5^, and are thought to be associated with evolution of anisogamy and with regulation of life cycle transitions^6^.

Research on the biology and evolution of sex chromosomes has primarily focused on mammals, birds, fish and *Drosophila* ^1,7^, and consequently studied the conventional XX/XY and ZW/ZZ diploid sex determination systems. In contrast, the U/V haploid sex-determination systems such as those in bryophytes and algae (brown, red and green lineages)^8,9^ have remained largely unexplored. In U/V systems, sexes are not determined at fertilization but instead during meiosis, when haploid spores receive a U chromosome, forming a female gametophyte, or a V chromosome, forming a male gametophyte^10^. These fundamental differences between U/V and the XX/XY and ZW/ZZ systems have broad evolutionary and genomic implications^11^. However, to date, only the U/V systems of the brown alga *Ectocarpus* and that of the bryophytes *Ceratodon*^12^*, Sphagnum*^13^, *Marchantia*^14^, and the U chromosome of *Syntrichia caninervis*^15^ have been fully sequenced and assembled. These studies provided important insights into the contrasting features of the bryophytes’ U/V chromosomes, such as their propensity to degenerate, and the acquisition of sex-linked genes as a consequence of genetic conflict between sexes^16^ (reviewed in ^17^). Similarly, studies in brown algae, liverworts and volvocine algae have elucidated several mechanisms leading to the loss of the U/V system towards the emergence of co-sexual individuals (monoicy)^18–21^. However, understanding the full evolutionary history of the green lineage U/V system has remained out of reach due to the fragmentary nature of the available data. In addition, these species are highly divergent (500 Mya), do not share homologous U/V chromosomes^22^, and some have gone through whole genome duplication events which poses challenges for comparative genomic analysis^23^. Thus, we lack a broad-scale view across several U/V sex chromosome systems that would inform a reconstruction of their evolutionary history.

In this context, brown algae (Phaeophyceae) represent exceptional models for investigating the origins and evolution of sex chromosomes. They span a bewildering variety of reproductive systems, life cycles and sex chromosome systems in a single lineage, a feature that is unique among Eukaryotes^24^. Moreover, separate sexes appear to be the ancestral state in brown algae^24^, suggesting that sex chromosomes and their dimorphic sexes share a common origin.

Here, we exploit a range of brown algal species and outgroups with diverse types of sexual systems to study the origin, evolution and demise of U/V sex chromosomes. By reconstructing the evolutionary history of U/V sex chromosomes in brown algae, our work opens up new research directions for the study of genomes and sex chromosome evolution in a large taxonomic context, and improves our biological understanding of this ecologically relevant eukaryotic lineage.

## RESULTS

### The origin of brown algal sex chromosomes

Among eukaryotic organisms, brown seaweeds span a rich variety of morphologies, life cycles and sexual systems^9,24,25^. Using high-quality reference genomes^26,27^, supplemented with genetic and HiC physical maps (see methods), we generated chromosome or near-chromosome level genome assemblies of nine representative brown algal species. Combined with the outgroup species *Schizocladia ischiensis*^28^, these span the phylogenetic, morphological and reproductive diversity of brown seaweeds^25^ (**Fig. 1A**). Overall, our analyses reveal that brown algae have a relatively stable karyotype (27-33 chromosomes) and a very well conserved macrosynteny (*i.e.*, similar blocks of genes located in the same relative positions in the genome across species) (**Fig. 1A**).

**Figure 1.**
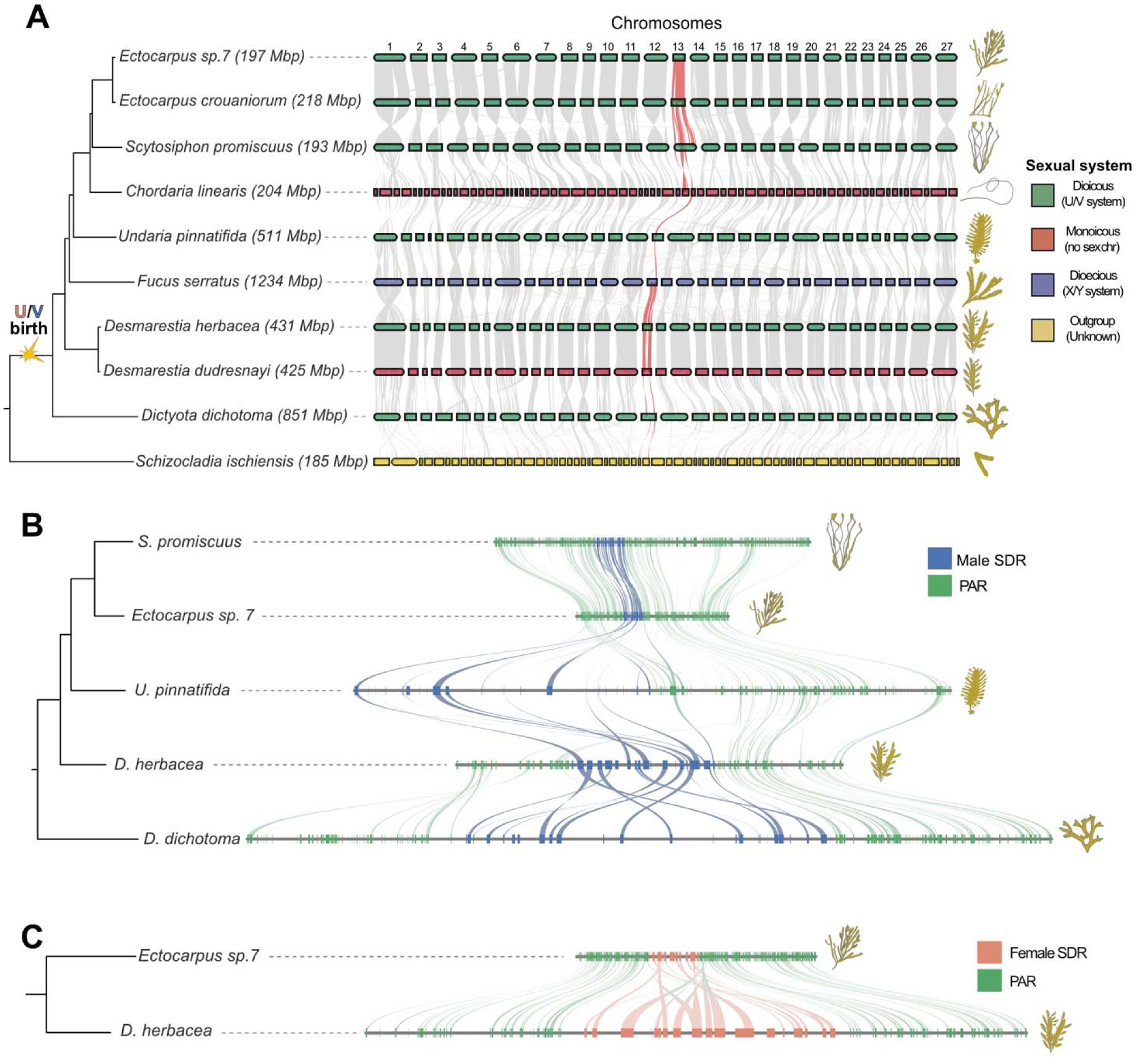
Origin of the brown algal U/V sex chromosomes. (A) Macrosynteny plot between brown algal genomes of six dioicous species (green), two monoicous species (red), one dioecious species (blue) and the outgroup species (yellow). The syntenic blocks corresponding to the V sex chromosome are highlighted in red, and the birth of the U/V chromosomes is displayed in the phylogeny. The size of each genome is displayed inside brackets. (B) Microsynteny plot of the V chromosomes in five dioicous species, showing the regions that belong to the male sex-determining region (blue) and the PAR (green) for each species. (C) Microsynteny plot of the U chromosomes in two dioicous species, showing the regions that belong to the female sex-determining region (peach) and the PARs (green) for both species.

By combining comparative genomic approaches with experimental validation of sex-linkage using PCR (see methods ‘Discovery of the UV sex determination regions’) we identified the female and male SDRs (U-SDR and V-SDR, respectively) on the sex chromosomes for each dioicous species (**Fig. 1B-C**). Consistent with dioicy being the ancestral state in the brown algal lineage^24^, the synteny analysis demonstrated that all species with a U/V sexual system share the same sex chromosome, including the early-diverging species *Dictyota dichotoma*. Based on these data, we conclude that the recombination suppression event that gave rise to the U/V sex chromosomes occurred within the branch that separates *Schizocladia* from *D. dichotoma* (**Fig. 1A**), between 450 Mya at the origin of the Phaeophyceae, and 224 Mya at the crown node of the SSD + BACR clade^29^. The master sex-determining gene *MIN*^30^ is present inside the V-SDR of every dioicous brown algal species, including the early diverged *D. dichotoma*. When analyzing other algal draft genomes^26^, we found that *MIN* was present in the most early-diverging brown algae species; *Discosporangium mesarthocarpum* (mRNA_D-mesarthrocarpus_Contig843.5.1), potentially pushing the latest date of the U/V system to the crown node of the brown algae (370 Mya^29^). However, it remains unknown whether the U/V system functions in sexual determination in *D. mesarthocarpum*. We note that neither *Fucus serratus* nor the monoicous (co-sexual) species *Chordaria linearis* and *Desmarestia dudresnayi* have a U/V sexual system, yet they still conserve *MIN* within a chromosome homologous to the U/V sex chromosome in the dioicous species (hereafter referred to as ‘sex-homolog’).

In the outgroup *S. ischiensis*, we found very low synteny with the sex-homolog and patterns of fusion-with-mixing^31^ between the homologs of chromosomes 4 and 9 and between chromosomes 23 and 24 of *D. dichotoma* (**Fig. 1A; Fig. S1**). Such fusion-with-mixing events are believed to represent irreversible states of chromosome evolution^31^, and therefore, *S. ischiensis* likely represents a divergent karyotype, rather than a proxy karyotype of the last brown algal common ancestor. As noticed previously^30^, the *MIN* ortholog is present in *S. ischiensis*, suggesting that *MIN* could have played a sex-determining role as early as the common ancestor of the Phaeophyceae and *Schizocladia*. Therefore, the presence of *MIN* in this distantly related organism could push the emergence of U/V system as far back as the crown node of *Schizocladia* and Phaeophyceae (450 Mya)^30^.However, similarly to *D. mesarthocarpum*, *S. ischiensis* only reproduces vegetatively in culture, precluding the investigation of its sexual system.

We next examined the SDRs on the U and V sex chromosomes by comparing male and female genomes for all dioicous species (see methods). The V-SDR differed markedly across species, both in terms of gene content, total size and relative size compared to the PAR region (**Fig. 1B, Table S1**). We found the smallest V-SDR sizes in the Ectocarpales (*S. promiscuus* and *Ectocarpus* sp.7) (**Fig. 1B-C**), and noticed that 22 and 19 genes that correspond to PAR genes in *Ectocarpus.* sp.7 and *S. promiscuus,* respectively, were present in the V-SDR of brown algae with larger V-SDRs such as *U. pinnatifida*, *D. dichotoma* and *D. herbacea*. Similarly, the size of the U-SDR is different in *Ectocarpus* sp. 7 and *D. herbacea,* where 11 PAR genes bordering the U-SDR in *Ectocarpus* sp. 7 were incorporated in the U-SDR of *D. herbacea* (**Fig. 1C**). In addition, the V-SDR of *D. herbacea* and *D. dichotoma* has undergone substantial rearrangements and inversions whereas the V-SDR in *U. pinnatifida* has relocated from the center to the periphery of the V chromosome (**Fig. 1B**). Furthermore, we identified a large inversion in the U-SDR of *Ectocarpus* sp. 7 and *D. herbacea* (**Fig. 1C**).

Taken together these results indicate that the U/V sex chromosomes of the brown algae emerged between 450 and 224 Mya, via an inversion that suppressed recombination in a locus that contained the master male-determining factor *MIN*. The presence of *MIN* in distantly related lineages could push the age of the U/V chromosomes further back in time, but more evidence would be required to establish that dioicy existed in these organisms.

### The evolution of the U/V-SDR is associated with changes in the level of sexual dimorphism

A general feature in diploid sex chromosome evolution is the existence of evolutionary strata. These represent different recombination suppression events over time and can be detected by plotting synonymous substitutions per site (*Ks*) across male and female gametologs^32^. However, as previously observed in bryophytes^17^, the detection of evolutionary strata is challenging in U/V systems due to the shuffling of gene order and the dynamic movement of genes in and out of the U/V-SDRs^33,34^. These events disrupt the expected co-linearity in the evolutionary strata and limit the ability to clearly define them solely based on gene position. We observed an inverted pattern between the V-SDR and the U-SDR in *Ectocarpus* sp. 7 and in *Desmarestia herbace*a (Fig. 2A**-B**), consistent with the hypothesis of inversions leading to suppressed recombination between sex chromosomes. Upon analysis of *Ks* values across male and female gametologs, we noticed that the U/V-SDRs of *Ectocarpus* sp. *7* are composed of several highly-divergent gametologs (Ks > 1) in two (presumably older) ‘evolutionary strata’ and two gametolog pairs in a more recent evolutionary stratum (*Ks* < 1; Fig. 2B**-C****, Table S2**). In contrast, the U/V-SDRs of *D. herbacea* contained fewer gametolog pairs with *Ks* values higher than 1 and a higher proportion of recently diverged U/V-SDR genes (*Ks* < 1), forming at least four ‘evolutionary strata’ (Fig. 2B**-C**). The gametolog *Ks* values are broadly concordant between the U/V-SDR orthologs in *Ectocarpus sp. 7* and *D. herbacea* (**Table S2**), indicating that the accumulation of synonymous substitutions can be considered a proxy of evolutionary distance in comparable species (Fig. 2C). The occurrence of recently evolved evolutionary strata in *D. herbacea* combined with an inverted pattern between male and female is consistent with an expansion of the U/V-SDRs in brown algae caused by nested inversion events that suppress recombination. Indeed, when comparing *Ectocarpus* sp. 7 and *E. crouaniorum*, we note that the V-SDRs are inverted with respect to each other (**Fig. S2**), indicating that inversions are persistent even between closely-related taxa.

**Figure 2.**
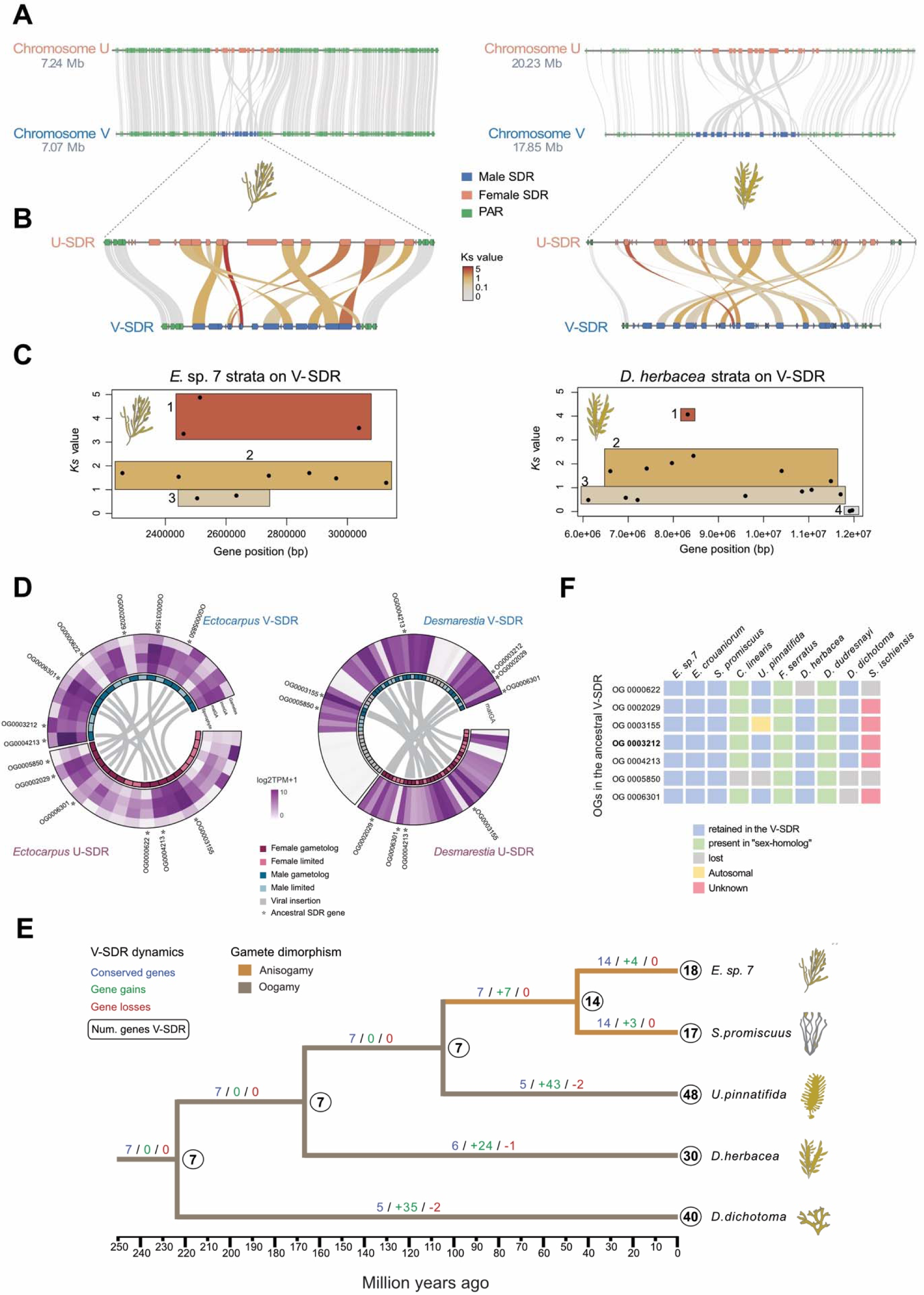
Lineage-specific expansion of the U/V-SDRs from a shared ancestral SDR is associated with increased sexual dimorphism. (A) Microsynteny plot between the U and V chromosomes of *Ectocarpus s p*an. d*7D. herbacea*. (B) Synteny between the U and V gametologs in the SDRs of *Ectocarpus sp. 7* and *D. herbacea*. The matching shades within the SDR are colored according to the number of synonymous substitutions per synonymous site (*Ks*) for each gametolog pair. (C) Identification of evolutionary strata in the male SDR of *Ectocarpus* sp. 7 and *D. herbacea* according to the *Ks* values of gametolog pairs and the position of the male (V) genes in the SDR. (D) Expression of SDR genes measured as log2(TPM+1) in *Ectocarpus* sp.7 and *D. herbacea*. For *Ectocarpus* sp. 7, expression levels are shown across different life cycle stages: gametes, juvenile gametophytes (immGA), sexually mature gametophytes (matGA), and sporophytes. For *D. herbacea*, data are available for sexually mature gametophytes (matGA). Gametologs are highlighted in dark colors (dark blue for V gametologs and dark purple for U gametologs), while sex-limited SDR genes are colored in light blue (V sex-limited) and light purple (U sex-limited). Viral insertions in *D. herbacea* V-SDR are marked in grey. Lines within the circos plot link corresponding gametologs in each species. Stars indicate the position of conserved SDR genes, which are also presented in panel F. (E) Ancestral state reconstruction of V-SDR gene content across the brown algal phylogeny, showing the expected number of genes in the SDR (circles on the right side of each node), the number of SDR genes that were retained compared to the previous node (blue), the genes that were incorporated into the SDR (green) and the genes that were lost or that were translocated outside of the SDR (red). The ancestral state reconstruction of gamete dimorphism (based on) shows a transition from oogamy to anisogamy in the Ectocarpales alongside with changes in the SDR gene content. (F) Schematic view of the location of the 7 ancestral V-SDR genes (blue: retained in the SDR; green: contained within the pseudoautosomal region or the sex-homolog; yellow: autosomal; red: Unknown genomic location; grey: lost). Bold: *MIN*. See also Table S4.

The presence of larger V-SDRs in early diverged brown algae such as *D. dichotoma* and smaller V-SDRs in later diverged orders such as Ectocarpales could indicate that gene loss may have reduced the physical size of the U/V-SDRs. However, different brown algal lineages could also have independently expanded their U/V-SDRs, while the U/V-SDR size of Ectocarpales reflect an ancestral state with fewer genes. To understand U/V-SDR size dynamics across species, we performed an ancestral state reconstruction of V-SDR gene content. Considering that the U-SDR and the V-SDR are expected to evolve similarly due to their inheritance patterns^33,35,36^, and overall better data quality for male genome assemblies, we focused our investigation on the V-chromosome. We found that V-SDR evolution is mostly driven by lineage-specific expansions of an ancestral V-SDR that contained a small number of genes in the common ancestor (Fig. 2E**-F**). Male gametologs with higher *Ks* values were mostly supported as ancestral V-SDR genes within the ancestral state reconstruction, whereas male gametologs that were independently acquired into the V-SDR displayed the lowest *Ks* values in *D. herbacea* (**Table S2,** Fig. 2C). However, we note that some gametologs did not follow this pattern, highlighting that defining evolutionary strata for brown algae based solely on *Ks* values is challenging.

The SDRs of *Ectocarpus sp. 7* and *D. herbacea* contain genes that are homologous between the U and V chromosomes, suggesting that these haplotypes descend from a common ancestral autosomal region (**Table S3**). Even though U/V-SDR gene numbers differ between species, they share a similar ratio of 53-61%% gametologs and 47-39%% sex-specific (*i.e.* genes only found on the male or female SDR; **Table S3**). In contrast, the V-SDR of *D. herbacea* contains a higher number of sex-specific genes, but this excess was driven by a recent insertion of viral sequences into the V-SDR (**Table S3**). Thus, genes present only in one sex were either acquired after the divergence of the U and the V SDRs or lost by the counterpart haplotype. Further-more, we found that the gene numbers within the U-SDR and the V-SDR coincide in both *Ectocarpus* sp. 7 and in *D. herbacea* (**Table S3**), further supporting parallel inheritance patterns for U and V chromosomes^33,35,36^.

Intriguingly, every estimation of V-SDR size (number of genes, total length and relative SDR-to-chromosome length) was strongly associated with extent of gamete sexual dimorphism for the five studied dioicous species (**Fig. S3A**). Specifically, oogamous species, in which male and female gamete size differs significantly (e.g., *U. pinnatifida* or *D. herbacea*) have larger V-SDRs than species with modest gamete size differences (anisogamy, e.g. *Ectocarpus sp. 7*) (Fig. 2E). In contrast, *Ectocarpus* sp. 7 and *Scytosiphon promiscuus* retain all ancestral V-SDR genes with few additional gene gains, indicating that the Ectocarpales resemble the ancestral state of the V-SDR more closely than the early-diverging lineages. Most of the V-SDR genes in the oogamous species were acquired autonomously for each lineage, with a single gene coding for an ATP-dependent RNA helicase (Ec-13_002220 in *Ectocarpus* sp. 7) acquired in the V-SDR of all oogamous species. Therefore, the expansions of the U/V-SDRs likely occurred independently and concomitant with changes in gamete size (Fig. 2E**; Fig. S3A**). Previous work suggested that oogamy is either ancestral in the brown algae^24^ or in the brown algal crown radiation clade^29^, and that a transition from oogamy towards anisogamy occurred in the Ectocarpales^24,29^. However, the independent expansion of the V-SDRs in each oogamous species and its association with increased gamete dimorphism (Fig. 2E**; Fig. S3A**) support an independent acquisition of oogamy from an isogamous or anisogamous ancestor. Thus, the association between increased gamete dimorphism and V-SDR gene content is either coincidental or the evolution of gamete sexual dimorphism is complex and cannot be explained solely by ancestral state reconstruction of life history traits^24,29^.

In our initial reconstruction, we retrieved 12 genes that were present in the ancestral V-SDR (see methods). However, our phylogenetic analyses revealed that five of those genes were present in several brown algal species as parallel and independent V-SDR expansions (**Fig. S3B**) and the ancestral V-SDR contained only seven genes (Fig. 2F). These seven ancestral V-SDR genes comprise six gametologs and one male-restricted gene, all possibly associated with aspects of sex determination (Fig. 2F**, Table S4**). For example, the master male sex-determining gene *MIN*^30^ is always present in the male SDR, consistent with its key role in sex determination. Another gene encoding a transmembrane protein with a putative sugar-binding and celladhesion domain (Ec-13_001840 in *Ectocarpus* sp.7), identified as a gametolog in the U-SDR (Ec-13_sdr_f_0020), is an interesting candidate for gamete recognition^37^. In addition, a STE20 serine/threonine kinase homolog (Ec-13_001910 in *Ectocarpus* sp.7) has a counterpart in yeast that is involved in the transcriptional activation cascade of mating-type genes after pheromone recognition^38^ whereas a casein kinase (Ec-13_001990), a MEMO-like domain protein (Ec-13_001810) and a GTPase activating protein (Ec-13_001710) represent signal transduction genes.

The six genes that belong to gametolog pairs showed no evidence of degenerative evolution and remained functional in both male and female *Ectocarpus* sp. 7. In *D. herbacea*, four genes are retained in both sexes, while the casein kinase was lost in the U-SDR and the putative carbohydrate-binding receptor was completely lost in both sexes, potentially indicative of a mild level of degradation.

We next examined the fate of genes from the PARs of *Ectocarpus* sp. 7 that became sex-linked following the expansion of the SDR in *D. herbacea*. Specifically, we observed two SDR expansions into the PAR1, a region containing four genes in *Ectocarpus* sp. 7, and into the PAR2, which contains 19 genes in *Ectocarpus* (**Table S5**). In the PAR2 region, six genes were identified as taxonomically restricted (TRGs) to the *Ectocarpus* lineage, while the remaining 13 genes are ancestral to all brown algae. Our analysis revealed that the majority of genes in the newly formed ‘strata’ in *D. herbacea* were retained as gametologs, with 11 gene pairs preserved. Two genes were retained as sex-limited genes, one on the U-SDR (mRNA_D-herbacea_F_contig86.15844) and one on the V-SDR (mRNA_D-herbacea_M_contig331.8898). One gene remained as a PAR gene in *D. herbacea*, and the final two were located on the *D. herbacea* autosomes. However, with only two species available for comparison, it is challenging to determine whether these genes moved into the PAR in *Ectocarpus* sp. 7 or moved out of the newly expanded SDR in *D. herbacea*. Together, these results suggest that despite being evolutionary old, haploid sex chromosomes in brown algae do not undergo substantial gene loss, and evolve mainly by gene gain, both from the duplication and translocation of autosomal genes as well as from the subsuming from neighboring PAR regions (Fig. 2F**; Table S5**).

The majority (58-100%) of U/V-SDR genes were expressed at substantial levels (log2(TPM+1) > 1) during the fertile gametophyte stage, consistent with purifying selection maintaining their biological activity (**Table S6**). Gametologs tended to have higher expression than sex-limited genes (Wilcoxon test, p-value=0.0007505 in *D. herbacea* and p-value=0.08843 in *Ectocarpus* sp. 7) (Fig.2D**, Table S6**). Furthermore, we conducted a comparative analysis of gene expression (mature gametophyte phase), in which we contrasted genes that became independently acquired in the SDR in each species with the single-copy autosomal paralogs present across different species. We found that the expression levels of the newly sex-linked genes remain comparable to their autosomal counterparts (**Fig. S4**). This observation suggests that the autosomal biological activity may either be co-opted to perform a male-specific function in the V-SDR genetic setting or be preserved due to its general importance for the gametophyte development. However, by examining expression across multiple tissues—such as gametes, immature gametophytes, and sporophytes, which is possible in *Ectocarpus* sp. 7 due to the availability of comprehensive data—we observed that the expression of U/V-SDR genes is not confined to mature gametophytes (Fig.2D). Together, our observation thus suggest that the U/V-SDR contains genes playing a broader role in development or more pleiotropic functions, in addition to more conserved genes with roles in sex determination and gametophyte fertility.

Whilst the size of the SDR was positively associated with the level of gamete dimorphism (see above), we found no correlation between the amount of autosomal sex-biased gene (SBG) expression and differences in gamete size across males and females (**Table S7**), in agreement with a study of a small subset of brown algal species^21^. Notably, species with low sexual dimorphism and smaller V-SDRs actually exhibited the highest proportion of SBGs (**Fig. S5A**). Intriguingly, we observed an enrichment of male-biased genes (MBG) within the PAR across all species, except in *D. dichotoma (***Fig. S5B**).

Most ancestral V-SDR genes were also present in the monoicous species *C. linearis* and *D. dudresnayi*, the dioecious species *F. serratus* and in the outgroup *S. ischiensis* (Fig. 2C**, Table S4)**, suggesting that they play important roles even in the absence of a U/V sex-chromosome system. All the non-dioicous brown algal species retain most of these genes on their sex-homolog chromosomes, as expected from the strong conserved synteny in Phaeophyceae. However, the SDR orthologs in *S. ischiensis* are scattered throughout the genome, possibly as a consequence of fusion-with-mixing events.

Altogether, our analyses show that the U/V-SDRs in brown algae are structurally dynamic, and evolved mainly by lineage-specific gene gain concomitant with increased sexual dimorphism, via frequent inversions. We detect a set of genes that has been conservatively sex-linked across the evolution of the dioicous brown algae, suggesting that these genes may have a key role in sex determination and/or differentiation.

### Structural features and evolutionary dynamics of brown algal UV sex chromosomes

Our chromosome-level assemblies allowed us to evaluate structural features that differentiated V-SDR and PAR regions from the rest of the genome. As expected for non-recombining regions^39^, exons were depleted in brown algal V-SDRs (**Table S8**) whereas repetitive elements were enriched (Fig. 3A**; Table S9**). Within the repetitive element classification, we found that ‘unclassified’ transposable elements in the sex chromosome were enriched in *S. promiscuus*, and to a lesser extent, in *Ectocarpus* sp. 7 (Fig. 3A**; Fig. S6; Table S10**). In contrast, the V-SDRs of species that underwent genome expansion (*U. pinnatifida*, *D. herbacea*, *D. dichotoma*) were colonized by LTR elements, which appear to be the main drivers of genome expansion in brown algae (Fig. 3A**; Fig. S6; Table S10**).

**Figure 3.**
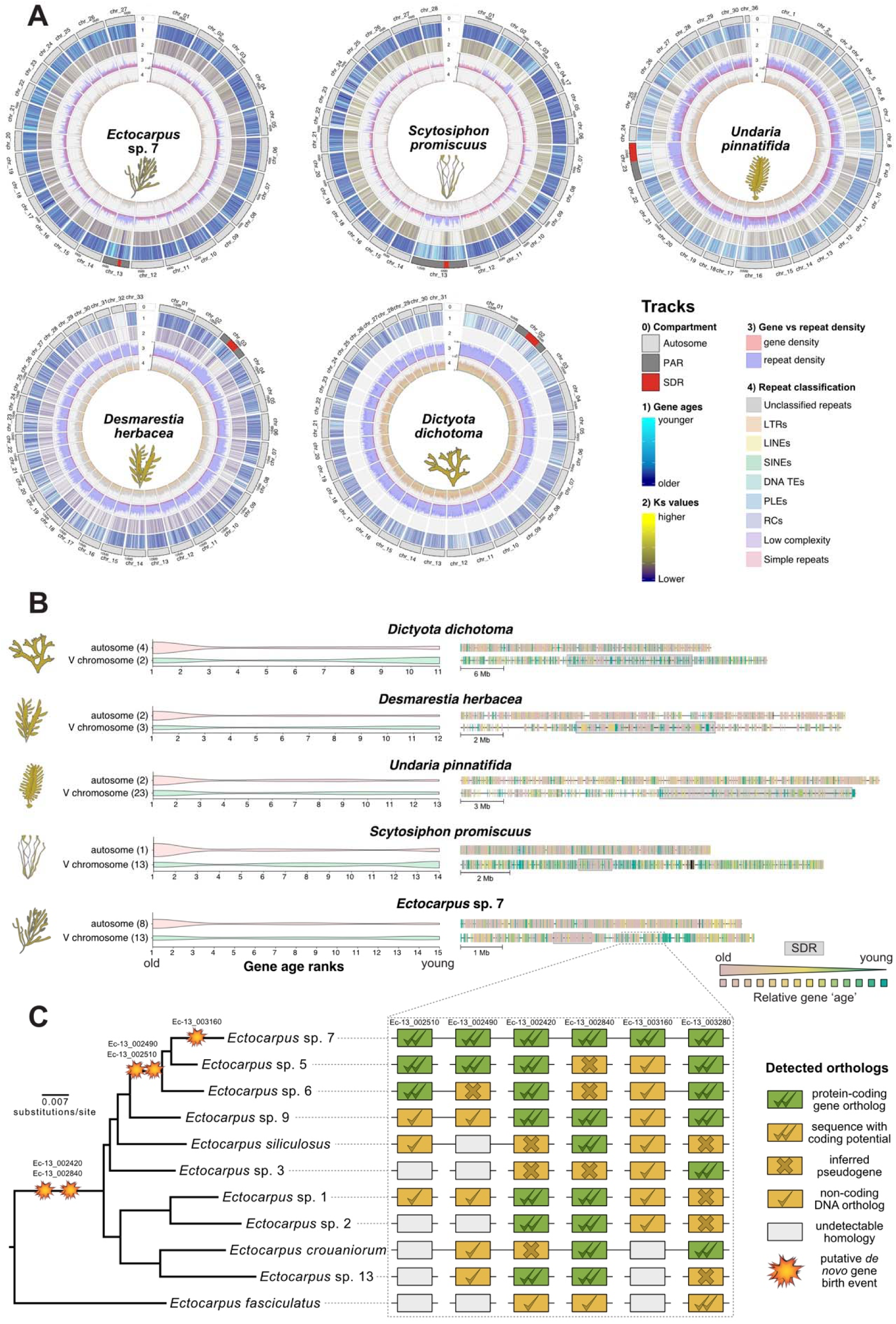
The U/V sex chromosomes act as cradles for new genes that evolve de novo. (A) Circos plots displaying tracks for the following genomic features: 0) chromosome: autosomes, PARs and SDR; 1) relative gene ages, 2) *Ks* values, 3) proportion of gene density (red) against repeat density (blue) and 4) repetitive element classification. (B) Violin plots and physical distribution of relative gene ages across one autosome and the V sex chromosomes of *Ectocarpus* sp. 7, S. *promiscuus*, *D. herbacea*, *U. pinnatifida* and *D. dichotoma*. The SDR of the V sex chromosomes are highlighted with a grey shadow. (C) Detection of putative *de novo* gene birth events in the PAR of *Ectocarpus* sp. 7 across the *Ectocarpus* phylogeny. The inferred status of pseudogene or noncoding DNA in the ortholog DNA region is based on ancestral state reconstruction using maximum likelihood.

We also detected fewer conserved sex chromosome orthologs between species (Chi-squared test, *p*-value < 10^-4^, **Table S11**), possibly reflecting increased numbers of evolutionary young TRGs, as we demonstrated in *Ectocarpus* sp.7^40^. When we performed phylostratigraphy analyses^41,42^ to evaluate the relative ages of each gene across brown algal taxa, we found that genes in the different age categories were not randomly distributed across chromosomes (Kruskal-Wallis test, p-values <10^-40^ for all dioicous species). Consistently, young genes were enriched whereas older, highly conserved genes displayed lower proportions in the sex chromosomes (Fig. 3B; **Table S12**). Even though all analyzed dioicous species share a common ancestor (either within the same order or broader taxonomic groups), the gene age categories that were enriched in the sex chromosomes included genes that are taxonomically restricted within the species, genus and family levels (**Figs. S7-S11**), suggesting that the pattern of TRG enrichment in the sex chromosomes arose independently in each lineage.

We previously proposed a model to explain how TRG accumulation through generation-antagonistic selection would favor retention of young sporophyte-beneficial loci in the sex chromosome of *Ectocarpus* sp. 7^40^. Here, we confirmed that sporophyte-biased genes were enriched in the sex chromosome of both *Ectocarpus* sp. 7 and the kelp *U. pinnatifida,* but less so in *D. dichotoma* and *S. promiscuus* (**Table S13).** Importantly however, the two latter species are unlikely to experience ‘generation-antagonism’ given that both generations are phenotypically similar in *D. dichotoma* and the sporophyte generation is significantly reduced compared to the gametophyte generation in *S. promiscuus*^21,24^.

TRGs can emerge from mutations in ancestral non-genic regions that either create coding potential through *de novo* birth processes^43^, through selection-driven sequence divergence that changes the gene beyond the point of recognition to other homologs (orthologs or paralogs)^44^ or via a constant accumulation of neutral mutations leading to failure of detecting homology^45^. To explore these possibilities, we first estimated the synonymous substitution rate (*Ks*) between orthologs in other closely-related species and searched for differences in *Ks* values between chromosomes and between different genomic compartments (V-SDR, PAR, autosomes). We reasoned that if synonymous mutations behave neutrally^46,47^, then the rate of synonymous substitution (*Ks*) can be employed as a proxy for the point mutation rate^48,49^. We found that V sex chromosomes for all dioicous brown algal species were associated with higher *Ks* values (Fig. 3A; **Table S14**), consistent with a higher mutation rate compared to autosomes. We were intrigued to see that the enrichment of young genes and higher *Ks* values appear to be localized in the PARs of both *Ectocarpus* sp. 7 and *S. promiscuus* (**Fig. S7-S8**), but noted that this pattern extended to the entire sex chromosome in species with large V-SDRs such as *U. pinnatifida*, *D. herbacea* and *D. dichotoma* (**Fig. S9-S11**).

To examine if the TRGs in the sex chromosomes represent putative *de novo* gene birth events (and not homology detection failure artifacts), we analyzed a subset of six genes contained within a region of the PAR in *Ectocarpus* sp. 7 displaying different degrees of conservation within the two youngest gene ages (genus and species-level genes). We compared these genes against the genomes of ten different *Ectocarpus* species^26^ in a phylogenetically-aware manner. We detected orthologous regions that either belonged to a protein-coding gene or to noncoding DNA in the other *Ectocarpus* species (**Table S15**). We then performed ancestral sequence reconstructions to infer the coding potential of the ancestral sequences and detect the enabling mutations behind the emergence of those genes. Through this approach, we showed that five out of the six analyzed genes represent potential *de novo* gene birth events^43,50^ from ancestral sequences lacking open reading frames, while one gene may represent either an ambiguous *de novo* gene birth event or a fast-evolving gene lacking structural conservation (Fig. 3C**; Fig. S12**). Additionally, we detected transcriptional activity (TPM > 1 at any life stage; **Table S13**) for four out of the five putative *de novo* genes, as well as for 82.3% of the total TRGs in the genome and 85.5% of the TRGs in the sex chromosome of *Ectocarpus* sp. 7 within the two youngest gene ages. This indicates that the TRG enrichment pattern in the sex chromosome is not driven by annotation artifacts, although these TRGs may also represent ORF-containing non-coding transcripts. Therefore, brown algal U/V sex chromosomes appear to act as ‘gene cradles’, fostering *de novo* birth of coding or non-coding loci.

Finally, in order to test the generality of this ‘gene cradle’ pattern in other UV systems, we used the same approach in other organisms with haploid sex determination systems and chromosome-level assemblies. Specifically, we analyzed *Ceratodon purpureum*, *Sphagnum angustifolium* and *Marchantia polymorpha*, three plant species that harbor U/V sex chromosomes^13,51,52^, and *Cryptococcus neoformans*, a fungal species that harbors mating-type chromosomes^53^. We observed clear gene cradle patterns in the V chromosomes of *C. purpureum* and *S. angustifolium* (**Figures S13-S14, Table S12**), whereas we could not find any discernible cradle pattern in the highly degenerated U/V chromosomes of *M. polymorpha* nor in the mating-type chromosome of *C. neoformans* (**Figures S15-S16, Table S12**).

Together, our analyses indicate that brown algal U/V chromosomes accumulate TEs and have decreased gene density, similar to non-recombining regions in other eukaryotes, and their higher *Ks* is suggestive of higher mutation rates. Remarkably, brown algal U/Vs are a hotspot for spawning gene novelty through putative *de novo* birth processes. This ‘gene cradle’ pattern extends beyond the brown algae, as it is also present in the *Ceratodon* and the *Sphagnum* U/V systems, indicating this may be a general feature of mildly degenerated haploid sex chromosomes.

### Fate of U/V sex chromosomes following transition to co-sexuality

Next, we investigated the evolutionary trajectory of brown algal genomes following the ‘loss’ of the U/V sex chromosome system. When we used our genomic datasets to assess the fate of sex chromosomes once they are in a co-sexuality context, we found that the large majority of genes in the sex-homologs of *Chordaria linearis* and *Desmarestia dudresnayi* are male-derived, indicating that monoicy in both species originated from a male genomic background (Fig. 4A**-B**). The sex-homolog of *C. linearis* contains several rearrangements that span both the regions belonging to the ancestral PARs and the ancestral SDR (Fig. 4A). Most of the U/V-SDR orthologs remained in a syntenic region within the sex-homolog, with only one V-SDR ortholog being translocated into another position within the same chromosome (ortholog of Ec-13_001930). Notably, most of the U/V-SDR orthologs that were lost during the transition to monoicy display their corresponding gametologs or retain closely-related autosomal paralogs within the genome of *C. linearis* (**Table S16**). Nonetheless, all these genes belong to multigene families in both *Ectocarpus* sp.7 and *C. linearis*. Therefore, it remains unclear if the expression of the lost U/V-SDR genes in *C. linearis* would be compensated by another gene family member.

**Figure 4.**
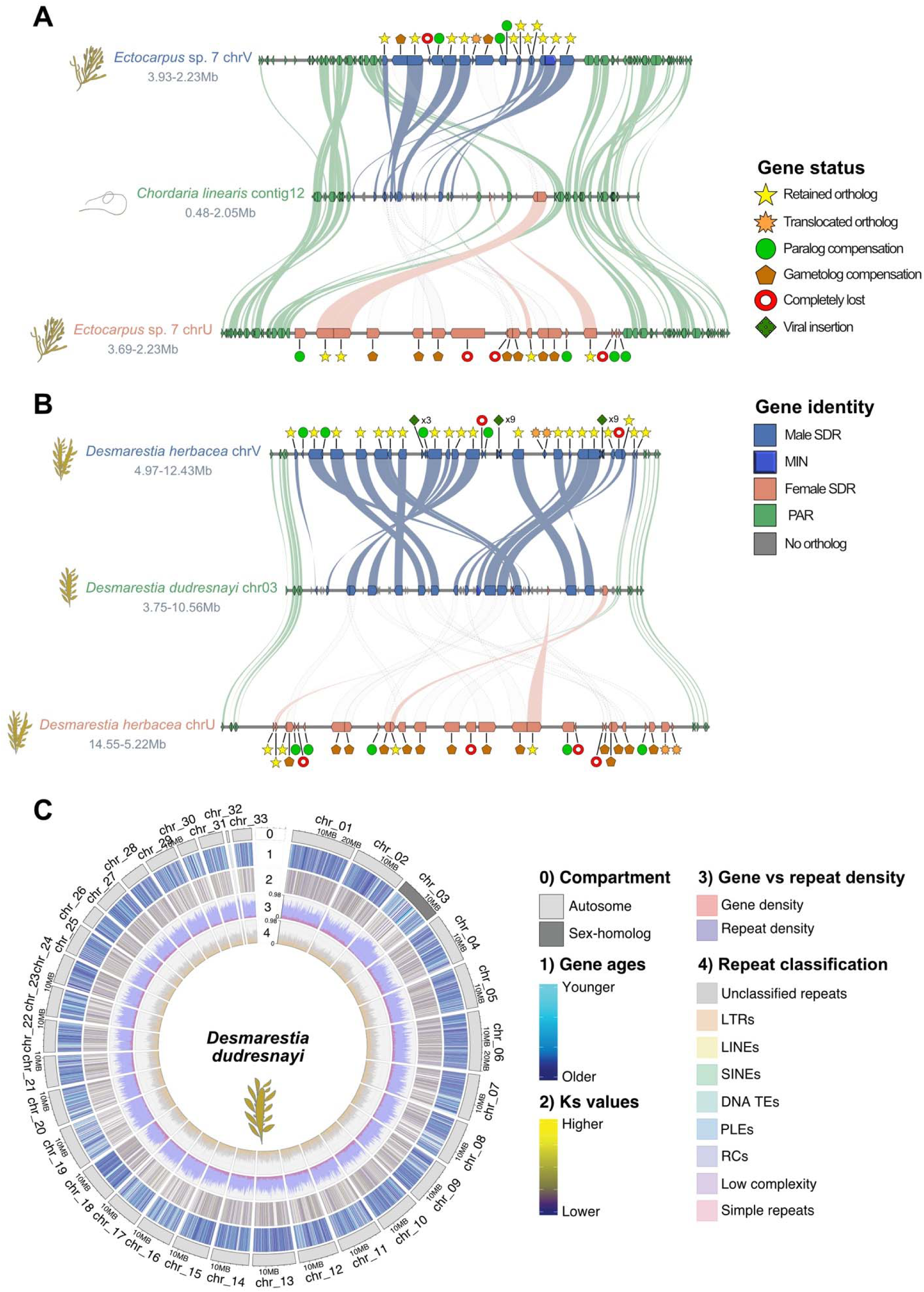
Fate of sex chromosomes during transitions from dioicy to co-sexuality. (A) Comparison of the sex-homolog in *C. linearis* against the U and V chromosomes of *Ectocarpus* sp. 7. (B) Comparison of the sex-homolog in *D. dudresnayi* against the U and V chromosomes of *D. herbacea*. The color code represents the identity of the genes alongside the chromosomes, while the shapes represent the evolutionary fate of each SDR gene in the monoicous genome. The matching shades between the SDRs and the sex homolog are either color-coded by their ancestral background or they appear as transparent dotted shades if the gametolog of the other sex was retained. (C) Circos plot of *D. dudresnayi* displaying tracks for the following genomic features: 0) chromosome compartments: autosomes and sex-homolog; 1) relative gene ages, 2) *Ks* values, 3) proportion of gene density (red) against repeat density (blue) and 4) repetitive element classification. *D. dudresnayi* retains the gene cradle pattern in the sex-homolog, but its *Ks* values are no longer different to those of the other chromosomes (see also Fig. S18 and Tables S8-S12, S14).

Interestingly, most of *C. linearis* V-SDR-derived genes are present in a contiguous region of the sex-homolog, while the three U-SDR-derived orthologs are framed by PAR orthologs that were translocated from one border of the SDR to the other and that separates the U-SDR insertions from the V-SDR-derived genes. Thus, *C. linearis* descended from a male genomic background that acquired its female-derived SDR genes through at least two non-homologous recombination events (Fig. 4A**; Fig. S17A**).

Similarly, the sex-homolog in *D. dudresnayi* also includes chromosomal rearrangements, with at least two inversion events within the region that is homologous to the V-SDR in *D. herbacea* (Fig. 4B), and with 20 V-SDR-derived orthologs and only five U-SDR-derived genes (**Table S17**). The two most recently acquired U/V-SDR genes in *D. herbacea* (stratum 4, Fig. 2B**-C**) are also present in the monoicous *D. dudresnayi*, but these two genes were integrated into the U/V-SDRs of *D. herbacea* after diverging from *D. dudresnayi*. Similarly, we found at least three viral insertions in the male SDR of *D. herbacea* that are absent in the sex-homolog of *D. dudresnayi*, suggesting that these insertions are male-exclusive and occurred after the two species had diverged. As with *C. linearis*, most of the genes that were lost during the transition to monoicy had a putative gametolog or a paralog compensation (**Table S17**). Three of the U-SDR-derived genes in *D. dudresnayi* are scattered throughout the ancestral V-SDR region, likely representing independent translocation events, with one of them appearing close to the PAR region in a similar fashion to *C. linearis*, while the other female genes were translocated and conserved elsewhere in the genome (Fig. 4B**; Fig. S17B**).

We examined next the expression pattern of ancestral male SDR genes in the sex-homologs of *C. linearis* and *D. dudresnayi*. Both retain a copy of *MIN*, confirming its crucial role in the male developmental pathway also in the monoicous species^30^. These genes are expressed during the reproductive stages of both species **(Table S18)**, suggesting that ancestral V-SDR genes play a key role in reproduction - even in the absence of a U/V sex-chromosome system. Interestingly, gene retention in all non-dioicous brown algal species occurs in a strongly conserved synteny pattern on sex-homolog chromosomes. Conversely, most of the U-SDR genes are absent in the monoicous species, with the exception of a putative intracellular cholesterol transporter (Ec-13_sdr_f_0007 in *Ectocarpus* sp.7) that is present in both monoicous species (**Table S18**) and is also conserved in the U-SDR of *D. herbacea* (mRNA_D-herbacea_F_contig563.13047.1). Additionally, this gene is actively expressed during fertility in both *Ectocarpus* sp. 7 and *D. herbacea* (**Table S6**), hinting to an important role in the female developmental pathway.

Together, these results indicate that monoicous species require a male genomic background to retain all necessary features of a functional sexual phenotype and likely arose from an ancestral male that acquired female SDR genes via ectopic recombination.

Lastly, we examined if the typical features of the UV sex chromosome such as high TE content, TRG enrichment and high *Ks* values remain present once the sex chromosomes do not perform their sex-chromosome function. We detected TE content enrichment and younger TRGs on the sex-homolog in *D. dudresnayi*, although the *Ks* values were not higher in this chromosome when compared to the rest of the genome (Fig. 4C**; Fig. S18; Tables S8-S12, S14**). Due to the recent loss of the U/V system in *D. dudresnayi*, the evolutionary vestiges of its past as a sex chromosome are still present in the sex-homolog, such as TRG enrichment and lower gene density, although the evolutionary patterns are gradually being lost, as shown by the *Ks* values.

### Transition from haploid to diploid sexual systems in the Fucales

Transition from haploid to diploid sex determination remains undescribed because in most eukaryotes the transition occurred in very deep evolutionary times precluding its examination. The Fucales are one of two brown algal lineages that recently transitioned to a diploid life cycle^54^, with many Fucales species having separate sexes in the diploid stage of the life cycle (dioecious sex determination^55^). Previous field observations indicate that *Fucus serratus* displays a sexual system with inferred male heterogamy^56^. Although we conducted high coverage genome sequencing of males and females, combined with RAD-seq data generated for 12 males and 12 females from a field population (see methods for details), we were unable to identify sex-linked sequences, suggesting that the SDR of *F. serratus* is likely to be very small and undifferentiated. Whilst the exact age of emergence of the X/Y system in *Fucus* is uncertain, ancestral state reconstruction analyses suggest that dioecy likely emerged in the last common ancestor of the *Fucus* genus^57^ between 25 and 5 Mya^26^. These data are consistent both with a young sex chromosome in *F. serratus*, having arisen following the transition to diploid life cycle^24^, and with the high turnover in the Fucales lineage^57^.

Our synteny analyses (see Fig. 1A) indicated that male *F. serratus* retains a chromosome that is homologous to the V chromosome of the ancestral U/V species. In addition, the conserved SDR genes from the ancestral U/V systems include the V-specific male-determining gene *MIN*^30,33^ and several other V-linked genes (see Fig. 2C), although some ex-SDR genes likely arose from the female gametolog (Fig. 5A). Although none of the ex-U/V-SDR genes are sex-linked in *F. serratus* males, *MIN* and four additional ancestral V-SDR genes have a strong male-biased expression pattern (Fig. 5A**, Table S19**) and were silenced in *F. serratus* females. As expected, these ancestral V-SDR genes were also silenced in female strains of three other Fucales species (*Ascophyllum nodosum*, *Fucus ceranoides* and *Fucus vesiculosus*; **Table S20**). Only one U-specific gene was found in *F. serratus*, and this gene was not differentially expressed between males and females (Fig. 5A) Thus, whilst the ancestral V-linked genes are no longer sex-linked in the Fucales, they are still likely participating in the genetic cascade of male sex determination and/or differentiation.

**Figure 5.**
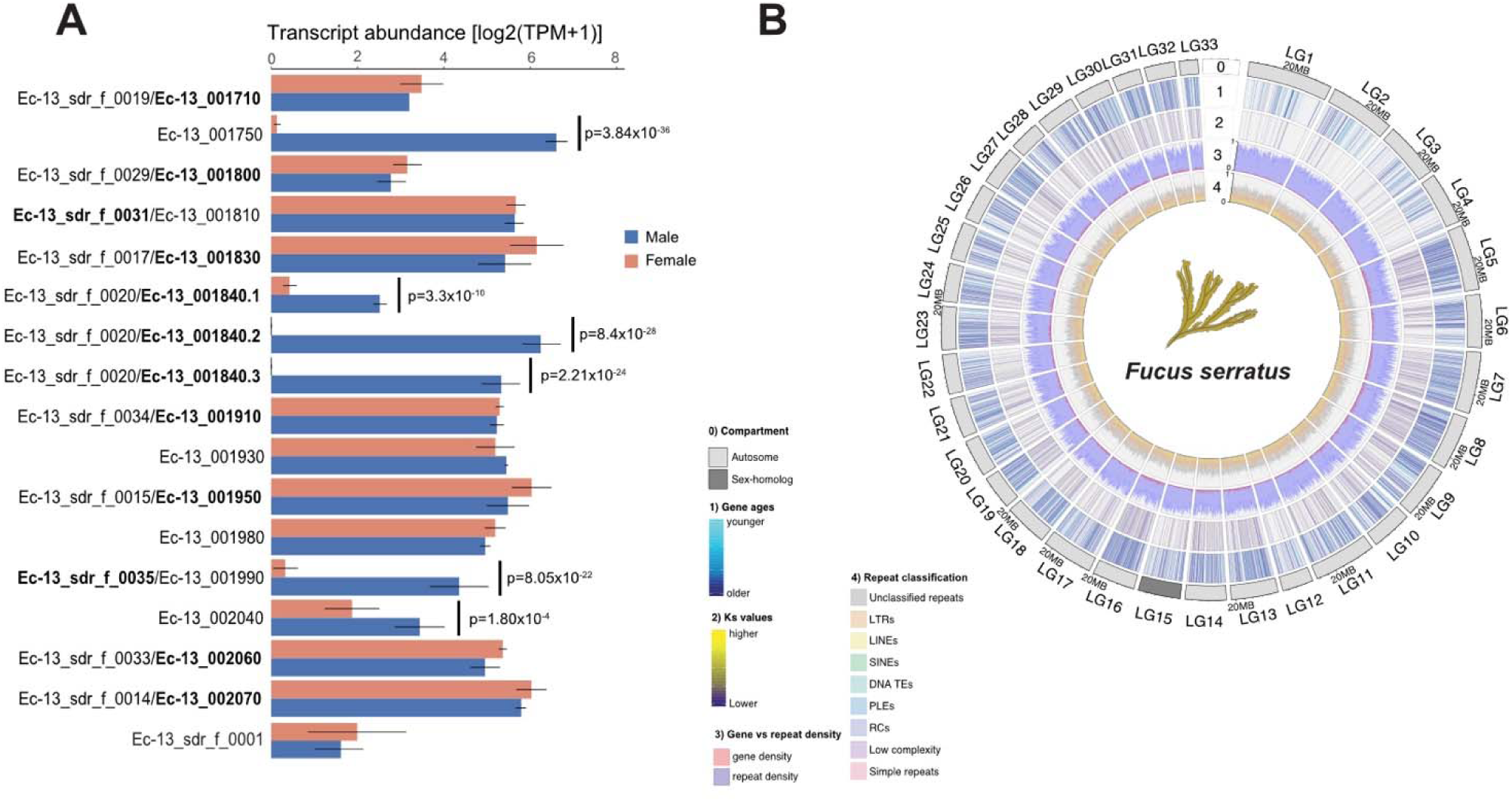
Transition from haploid to diploid sex determination. (A) Expression of ancestral U/V-SDR genes in the diploid species *F. serratus*. Gene expression of mature algae (using 3 males and 3 females, see methods) is given as log2(TPM+1) and bars represent standard deviation of the mean. Bold text represents whether the gene in *F. serratus* corresponds to an ancestral male or the female gametolog. (B) Circos plot of *F. serratus* displaying tracks for the following genomic features: 0) chromosome compartments: autosomes and sex-homolog; 1) relative gene ages, 2) *Ks* values, 3) proportion of gene density (red) against repeat density (blue) and 4) repetitive element classification. The sex-homolog has lost the genomic footprints of a sex chromosome, with no distinguishing patterns in its sex-homolog or in any other chromosome (see Fig. S19 and Tables S8-S12, S14).

Consistent with the lack of a differentiated sex chromosome, we did not observe any specific structural features in any of the chromosomes of *F. serratus*. In particular, we did not observe any trace of gene cradle pattern in *Fucus* chromosomes (as described above for U/V systems) nor higher *Ks* values or higher TE content on the sex-homolog or in any other chromosome (Fig.5B**; Fig. S19; Tables S8-S12, S14**).

## DISCUSSION

### The rise of brown algal sex chromosomes

Here, we characterized features of sex chromosomes or sex-homologs in nine representative brown algal species and one outgroup that originated between 450 and 244 million years ago^29^. Across species, we document an overall high degree of synteny conservation, similar to the evolution of multicellularity in Metazoans^58^. The ancestral cassette with seven V-SDR genes is highly conserved across all brown algal species, particularly those with dioicous sex determination, suggesting that a chromosomal inversion led to the joint sequestration of these genes into the non-recombining region during the birth of the U/V-SDRs. Given the pivotal role of *MIN* in brown algal male sex determination^30^ and its conservation across brown algae, this gene is the main driver behind the initial event that gave rise to the U/V system. Conversely, the other six genes in the ancestral V-SDR share gametolog pairs with the U-SDR, and their function could be relevant for both male and female reproduction. Whether their presence within the U/V-SDRs is essential to maintain the U/V system or whether they were sequestered accidentally during the initial recombination-suppression event remains elusive.

Despite conserved synteny in brown algal autosomes, the structure of the U/V sex chromosomes, and particularly the SDR, is highly dynamic. Frequent chromosomal inversions^59^ played a pivotal role in the evolution of brown algal U/V-SDRs. Inversions were involved both in the initial recombination suppression in proto-sex chromosomes, as well as subsequent expansion events of the U/V-SDRs into neighboring PARs. As described in other haploid systems^39,60^, TEs substantially accumulated in these non-recombining regions as a consequence of the initial recombination suppression in the SDR^61^, and likely led to further rearrangements via TE-mediated inversions.

### Independent SDR expansions are linked to sexual dimorphism and rise of morphological complexity

Models of XX/XY and ZW/ZZ sex chromosome evolution seek to explain how sexual dimorphism may drive SDR expansion in a way that favors evolution of new evolutionary strata^62,63^. We found that higher sexual dimorphism in gametes is associated with expanded U/V-SDR boundaries in brown algae, consistent with involvement of a larger number of genes under sexually antagonistic selection^64^. This relationship may reflect interplay between morphological complexity, the retention and acquisition of sex-specific genes in the SDR, and the emergence of sexual dimorphic traits. Although sexual antagonism is assumed to drive movement of genes into the SDR, the fact that newly acquired SDR genes show similar expression patterns to their autosomal orthologs in other species suggests that other mechanisms are at play in this process. Further analyses will be needed to demonstrate that the genes taking up residence in the U/V-SDRs are specifically important for reproductive functions rather than for the (vegetative) development of gametophytes.

Oogamy is thought to be an ancestral state in the early branches of the brown algal tree, but this trait seems to be highly labile^24,29^. Our analysis is concordant with a scenario in which several transitions to oogamy occurred independently in different lineages from a less dimorphic ancestor, and that the increase in gamete dimorphism was accompanied by modest expansion of the sex locus. Similar to the liverwort *Marchantia,* a streamlined gene content of the U and V SDRs in the brown algae could be regulating the autosomal gene network that controls sexual dimorphism. This contrasts with *Ceratodon*, where sex chromosomes harbor thousands of genes, and the number of sex-biased autosomal genes is minimal compared to sex-linked genes^51^. In support of this hypothesis, we detected substantial sex-biased gene expression in the mature gametophytic stages, which are potentially under the control of sex-linked genes. However, consistent with studies in other brown algae^21^ and in plants^65^, no correlation between levels of sex-biased expression and sexual dimorphism was observed. Thus, it is possible that the sexually dimorphic traits of the gametophyte are controlled by a subset of genes, while the majority of the sex-biased genes are instead relevant for gametophyte physiology or vegetative growth.

Established models of XX/XY and ZW/ZZ systems posit that the suppression of recombination between sex chromosomes is followed by accumulation of mutations and degeneration of the heterozygous chromosome (Y or W)^66^, whereas more recent models propose alternative mechanisms^67–69^. Our analyses reveal important differences in XX/XY and ZW/ZZ evolutionary trajectories, some of them previously observed in bryophytes^17^. The U/V structure in brown algae changes constantly through serial inversion events, TE accumulation and reduction of gene density. TE accumulation can be explained by suppression of SDR recombination, but the low gene density is not necessarily related to degenerative evolution and gene loss. On the contrary, our ancestral state reconstruction detected rare losses in the V-SDR across the brown algal phylogeny. The low gene density cannot be explained by gene movements out of the SDR either, since our ancestral state reconstruction coded any movement out of the V-SDR as a loss (such as gene Upin_00027965 in *Undaria pinnatifida*, see **Table S4**). Rather, the U-V-SDRs most likely expanded by an accumulation of TEs in noncoding regions, lowering the overall gene density.

Unlike the Y and W chromosomes, gene loss events in the U/V SDRs are rare. We found that *Ectocarpus* sp. 7 retained gametolog pairs for all six ancestral V-SDR genes, while *D. herbacea* retained most of these genes as gametologs, indicative of mild degeneration, as previously reported^3516,51^. In contrast with XX/XY and ZW/ZZ systems, our ancestral state reconstruction analysis documented more gene gain events than gene losses or gene movements outside of the V-SDR into the autosomes in all brown algal lineages. Considering that sex-determination occurs in the haploid stage, deleterious mutations are more efficiently purged than in diploid systems^70,71^. In addition, TE accumulation considerably differs in U/V sex chromosomes relative to autosomes. *Scytosiphon promiscuus* and *Ectocarpus* sp. 7, with the smallest genome sizes, display an enrichment of unclassified TEs in both SDR and PAR regions, whereas species with larger genomes no longer exhibit this pattern, and instead feature higher densities of LTR retroelements throughout the genome. While retrotransposons have an unbiased insertion pattern throughout the genome, DNA transposons display local hopping, meaning that they tend to insert themselves in the vicinity of the donor locus^72^. DNA transposons are overrepresented among unclassified repeats^73^, and these unclassified TEs may correspond to DNA transposons that flourished in the SDR and subsequently expanded to the PARs through local hopping. This signal would be subsequently diluted in species with larger genomes, as LTR elements increasingly colonized the genome, with a higher density in the SDR due to suppressed recombination.

### U/V sex chromosomes are gene nurseries

Brown algae U/V sex chromosomes exhibit an excess of TRGs, and our analysis suggest that a subset of these TGRs may have evolved *de novo* from previously non-coding regions, either as novel protein-coding ORFs or as non-coding transcripts. This led us to propose that U/V sex chromosomes function as ‘gene cradles’, likely through a combination of fast sequence divergence and *de novo* gene birth. What could be the mechanism underlying the U/V sex chromosomes ‘cradle’ pattern? Sex chromosomes in brown algae are enriched in repression-associated (hetero)chromatin marks (*e.g.*, H3K79me2)^74^, likely involved in suppressing TEs in this chromosome^75^. Heterochromatic regions have been suggested to display higher mutation rates due to the limited access of the DNA repair machinery to correct errors during replication^76^. Accordingly, we consistently documented higher synonymous substitution rates in the sex chromosome, which could be interpreted as higher mutation rates that would, thus, enable the emergence of young TRGs. Alternatively, the high density of DNA transposons within the U/V sex chromosomes could promote the co-option of their regulatory motifs and enable *de novo* birth of new transcripts, as has been described in *Drosophila*^77^. In either case, these TRGs could then be retained in the sex chromosome by ‘generation-antagonistic selection’, which would selectively maintain genes that bring an advantage to the sporophyte generation^40^. This mechanism requires differential selective pressures between the gametophyte and the sporophyte stages, and would thus require a dimorphic life cycle where gametophyte and sporophyte generations have considerable phenotypic differences. Notably, *D. dichotoma* does not display generation dimorphism^24^, and, accordingly, does not display a clear pattern of sporophyte-biased gene expression enrichment in the sex chromosome. Furthermore, we found no evidence for sporophyte-biased gene expression in the sex chromosome of *S. promiscuus,* which has a highly reduced sporophyte generation (in terms of size and morphological complexity)^24^ where selective pressures should be limited. Thus, the gene cradle pattern could be reinforced through generation-antagonistic selection, but DNA transposons and higher mutation rates alone may be sufficient to initiate this pattern in species lacking sporophyte-biased gene expression.

The gene cradle pattern is only present in brown algal species with U/V sex chromosomes, and is considerably diluted in *D. dudresnayi* where the transition to co-sexuality, and corresponding loss of sex chromosomes, occurred not later than 10 Mya when *D. dudresnayi* and *D. herbacea* diverged^26^. Moreover, it is absent in the dioecious *F. serratus*, that diverged from the ancestral U/V brown algae from 112 to 65 Mya^29^. These findings strongly indicate that the gene cradle pattern fades when the U/V system is lost.

How general is the gene cradle pattern outside brown algae? We could not detect enrichment of TRGs on small and degenerated U/V sex chromosomes such as those of *Marchantia polymorpha* ^52,78^. We note also that mating-type chromosomes such as those of *Cryptococcus neoformans*^53^ do not have a gene cradle pattern. In contrast, haploid chromosomes of *C. purpureus* ^16,51^ and *Sphagnum angustifolium* ^13^ exhibit a gene cradle pattern. The gene cradle pattern appears thus to be specific to U/V systems where chromosomal degeneration is mild and linked to haploid-diploid life cycles where the sporophyte stage is sufficiently complex, highlighting a key role for generation-antagonistic selection^40^. Therefore, our study reveals an interplay between complex life cycles, heterochromatic landscape, presence of DNA transposons, and higher mutation rates that could favor *de novo* gene birth processes, and emphasizes how common mechanisms may be at play across distant eukaryotic kingdoms.

### The demise of U/V systems

Monoicous brown algae were previously reported to display a transcriptomic profile that is closer to that of ancestral females^21^. However, our results clearly show that co-sexuality in *C. linearis* and *D. dudresnayi* arose from a male ancestor that acquired female genes. Generally, the expression of male-biased genes tends to be tissue-specific while female-biased genes tend to be broadly expressed^79^. This may explain the similarity of the transcriptional profiles between females and hermaphrodites in the brown algae^21,79^. The male developmental pathway may require more elements from the V-SDR for its proper activation, such as the master determinant factor *MIN*^30,33,80^, which would explain the convergent transition to monoicy from a male background. The transition to co-sexuality through a male genetic background has also been demonstrated in the liverwort *Ricciocarpos natans*^20^ and the volvocine alga *Pledorina starrii*^19^. The bisexual individuals of *P. starrii* harbor the male SDR and only differ from the male individuals by harboring an autosomal bisexual factor^19^. In *R. natans*, an aneuploid spore containing both U and V chromosomes led to a male V chromosome was inherited largely intact, while essential U genes were distributed to several autosomes^20^. However, the two monoicous brown algal species we studied contain only a few female genes that were directly translocated to the V chromosome. Importantly, we identified a female gene that is consistently present in both monoicous species, as well as in the U-SDR of *Ectocarpus* sp. 7 and *D. herbacea*, suggesting that this gene may be relevant for the determination of the female developmental pattern in the brown algae. We propose that the transition to co-sexuality in brown algae occurred through ectopic recombination, which added essential female genes to the male chromosome. However, while the combination of essential female and male genes is required for a successful transition to co-sexuality, which sex chromosome is retained may be of lesser importance. For example, the transition to monoicy in *Volvox africanus*involved the retention of a region resembling the female SDR of the dioecious ancestor, while most genes from the male SDR are absent and only a multicopy array of the male-determining gene *MID* was found in different genomic locations^18^. These findings provide a foundation for understanding how genetic networks controlling sex determination can be subtly rewired to produce significant changes in UV sexual systems. Essentially, an event combining crucial U and V genes is necessary for the transition to monoicy, whereas the mechanisms and genetic background may vary between systems.

The evolution of the XX/XY system in F*. serratus* is associated with a transition towards a fully diploid life cycle in Fucales^24,57^. The transition from a U/V towards a diploid sexual system likely involved a stage of epigenetic sex determination^3,57^, implying that the XX/XY system in the *Fucus* genus is fairly young compared to the 180-million-year-old Y chromosome in mammals or the 140-million-year-old W chromosome in birds^81^. A small and undifferentiated Y-specific region could explain our inability to detect the sex chromosome in *F. serratus*. Nonetheless, we found all ancestral V-SDR genes in the sex-homolog of *Fucus*, several of which displayed a male-biased gene expression across several Fucales species, particularly *MIN* that acts as the master male-sex determining factor in the U/V system^30^. Our results indicate that *MIN* and possibly other ancestral V-SDR genes are still involved in the male differentiation pathway, but likely changed their position downwards into the sex differentiation cascade. These results support and extend the “bottom-up” hypothesis of sex determination, where downstream components of sex differentiation are evolutionarily conserved between taxa, and where master-sex determining genes can be pushed downwards in the cascade and be replaced by new master regulators^82^. Together, our data are consistent with a step-wise transition to dioicy involving monoicy, and not directly from a UV to a diploid sex determination.

## CONCLUDING REMARKS

Here, we reconstructed the evolutionary history of the brown algal U/V system and formulate a hypothetical model for the evolution of haploid sex chromosomes (Figure 6). The U/V chromosomes of the brown algae descend from an ancestral autosome that contained *MIN* and other important genes that would later represent the ancestral SDR. This region underwent an inversion event that initiated the differentiation of the SDR between the U and V chromosomes. The SDR subsequently expanded through the accumulation of TEs and nested inversions in each algal lineage, engulfing genes that were previously contained in the PARs. The increase in size of the SDR was further caused by acquisition of genes from autosomes. It is possible that these autosomal genes were under sexual antagonistic selection and conflict would be solved by full sex linkage of these loci, but for a subset of these genes the role in sex was acquired only when they entered the SDR. Expansions of the SDR were, accordingly, positively associated with increased sexual dimorphism as each brown algal lineage evolved, and genes relevant for sex were retained within the SDR. It is possible that the continuous expansion of TEs increased the propensity of sex chromosomes to co-opt transcriptional motifs from DNA transposons that enabled new intergenic sequences to gain transcriptional activity. This TE expansion was likely limited by the recruitment of repressive chromatin marks and subsequent heterochromatinization, which would indirectly decrease the levels of DNA repair in the sex chromosome and further facilitate the emergence of young TRGs, some of them putative *de novo* genes or noncoding-transcripts, that would be then maintained through generation-antagonistic selection. The demise of the U/V sex chromosomes occurs in species that switch towards monoicy or to XX/XY. Transition to monoicy arose via introgression of female SDR genes in a haploid male genetic background. In the derived XX/XY system, the conserved sex-determining gene (*MIN*) from the ancestral V sex chromosome is no longer the master male-determinant, and has moved down the sex-determination hierarchy. Finally, ex-sex chromosomes gradually lose the genomic and evolutionary footprints of their past life as U/V sex chromosomes, such as the accumulation of TEs and the enrichment of young TRGs, as recombination resumes in these genomic regions.

**Figure 6.**
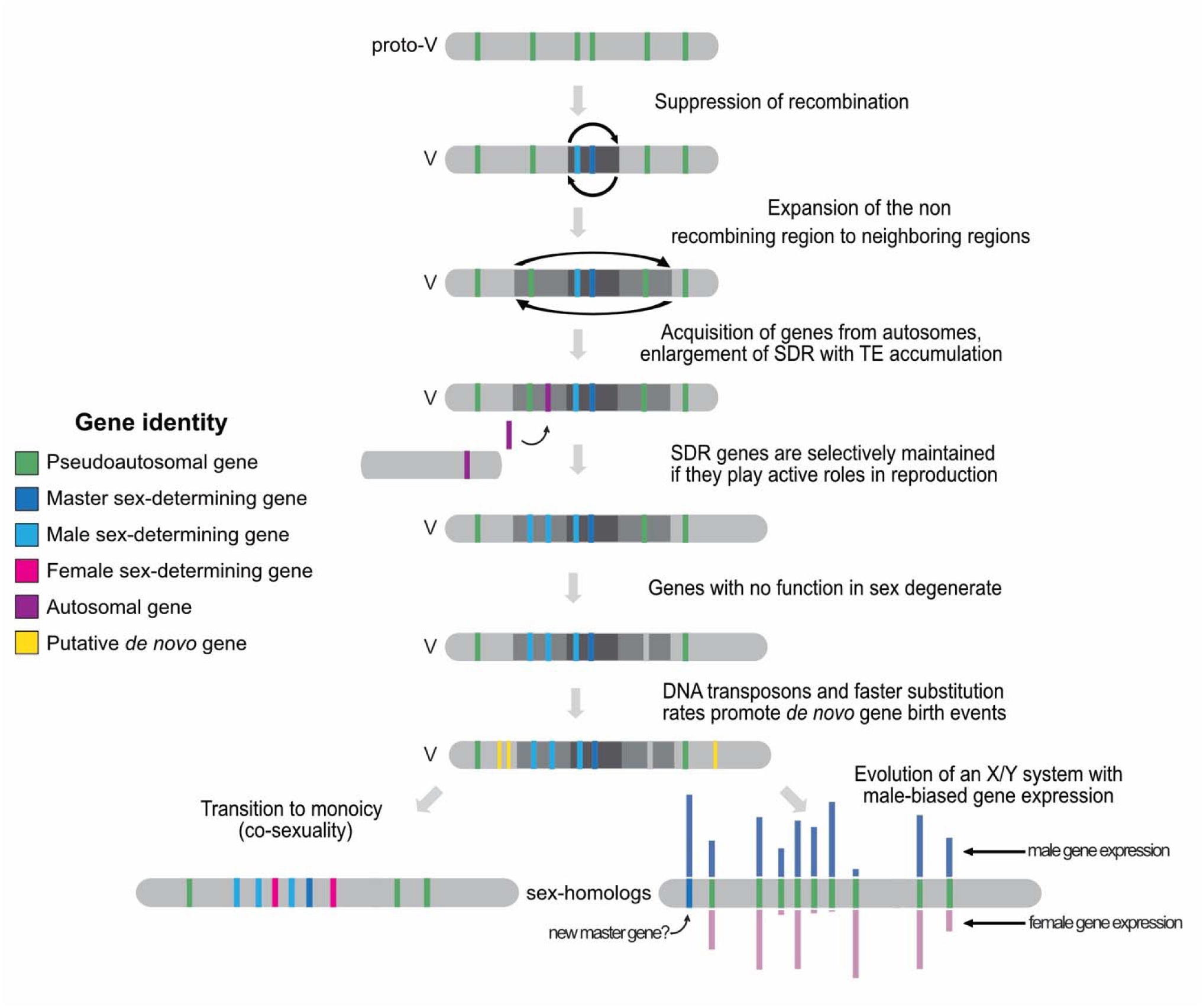
Hypothetical model for U/V sex chromosome evolution. U/V sex chromosomes arose from an ancestral autosome, via suppression of recombination that likely occurred via an inversion. The SDR expanded into neighboring pseudoautosomal regions (PAR) via inversions, but also by recruitment of genes from autosomes; expansion occurred in a lineage-specific fashion, concomitant with increased sexual dimorphism of the different species. SDR genes are maintained within the SDR if they have roles in sex, whereas genes with no role in sex are lost. Faster substitution rates, likely driven by the heterochromatic context of the sex chromosome may promote the rise of young genes, which are selectively maintained on the sex chromosome if they have advantages to the sporophyte generation. In species that switch to a diploid life cycle, the U/V system disappears, but the genes that are in the V-specific region retain roles in sex, although they are no longer masters. Transition from U/V separate sexes to co-sexuality (hermaphroditism) occurred when a male haploid individual acquired female-specific genes via ectopic recombination. During the demise of the U/V sex chromosomes, their structural and evolutionary footprints slowly erase.

## ACKNOWLEDGEMENTS

This work was supported by the MPG, the CNRS, Sorbonne University, the ERC (grant n. 864038 and 638240 to SMC), the France Génomique National infrastructure project Phaeoexplorer (ANR-10-INBS-09), the JSPS Overseas Research Fellowships (to MH), the BMBF-funded de.NBI Cloud within the German Network for Bioinformatics Infrastructure (de.NBI) (031A532B, 031A533A, 031A533B, 031A534A, 031A535A, 031A537A, 031A537B, 031A537C, 031A537D, 031A538A), the Investissements d’Avenir project Idealg (ANR10-BTBR-04-01), the European BG-01 BlueGrowth H2020 project Genialg (727892) and the ANR project Epicycle (ANR-19-CE20-0028-01). SMC is supported by the Moore Foundation (GBMF11489) and the Betten-court-Schuller Foundation. JBR is supported by a Humboldt Research Fellowship for postdoctoral researchers from the Alexander von Humboldt Foundation. We thank the members of the Phaeoexplorer consortium, in particular Chloe Jolivet, Leticia Mest and Delphine Scornet for assistance with algae cultures, Corinne Cruaud for help with sequencing libraries preparation, Erwan Corre and Arthur Le Bars for support with the Phaeoexplorer database and Arnaud Couloux for the genome assemblies and annotations. We are grateful to the Roscoff Bioinformatics platform ABiMS (http://abims.sb-roscoff.fr), part of the Institut Français de Bioinformatique (ANR-11-INBS-0013) and BioGenouest network, for providing computing and storage resources.

## AUTHOR CONTRIBUTIONS

JBR, APL: Investigation (equal); Formal analysis (equal); Methodology (equal); Visualization (equal), Writing – original draft (equal)

PL: Investigation (supporting); Formal analysis (supporting)

ED, GC, OG, KB, MH, KA, GL, EA, DL, RL, OG, SH, ZN, LG, AFP: Investigation (supporting) GH, JMA, GP, PW, FD, JMC: Data curation (supporting); Data acquisition (supporting)

FBH: Investigation (supporting); Methodology (equal); Data curation (equal); Formal analysis (supporting)

SMC: Conceptualization (lead); Funding acquisition (lead); Methodology (equal); Project administration (lead); Supervision (lead); Visualization (supporting); Writing – original draft (equal); Writing – review and editing (lead).

## DECLARATION OF INTEREST

The authors declare no competing interests

## SUPPLEMENTAL INFORMATION

**Table S1.** General characteristics of the genomes and UV sex chromosomes in five dioicous brown algal species.

**Table S2.** *Ks* values and evolutionary strata of the V-SDR gametologs of *Ectocarpus* sp. 7 and *Desmarestia herbacea*.

**Table S3.** Gametologs and sex-specific genes for the male and female SDR in *Ectocarpus* sp. 7 and *Desmarestia herbacea*.

**Table S4.** Ortholog table of the 7 ancestral male SDR genes in 10 brown algal species.

**TableS5.** SDR expansion into the PAR in *Desmarestia herbacea* and the fate of the newly sex-linked genes.

**TableS6.** Expression (log2(TPM+1) of male and female SDR genes during fertile gametophyte stage.

**Table S7.** Sex-biased gene expression between mature male and female gametophytes per species using DESeq2.

**Table S8.** FDR-corrected p-values for the pairwise Wilcoxon rank sum tests used to evaluate differences in the distribution of protein-coding gene content in 100kb windows across the chromosomes of seven brown algal species.

**Table S9.** FDR-corrected p-values for the pairwise Wilcoxon rank sum tests used to evaluate differences in the distribution of repetitive genomic elements in 100kb windows across the chromosomes of seven brown algal species.

**Table S10.** FDR-corrected p-values for the pairwise Wilcoxon rank sum tests used to evaluate differences in the percentage of unclassified transposons relative to the totality of repetitive elements in 100kb windows across the chromosomes of seven brown algal species.

**Table S11.** Chi-square test between the observed and expected value of orthologs for each chromosome in seven brown algal species.

**Table S12.** FDR-corrected p-values for the pairwise Wilcoxon rank sum tests used to evaluate differences in the distribution of gene age categories across the chromosomes of seven brown algal species, three plant species and one fungal species.

**Table S13.** Gene expression measured as log2(TPM+1) in gametophytes and sporophytes of different algal species.

**Table S14.** FDR-corrected p-values for the pairwise Wilcoxon rank sum tests used to evaluate differences in the distribution of synonymous substitutions per site (*Ks*) across the chromosomes of six brown algal species.

**Table S15.** Detection of orthologous sequences for six *Ectocarpus de novo* candidate genes.

**Table S16.** Evolutionary “fate” of the *Ectocarpus* sp. 7 sex-determining genes in *Chordaria linearis*.

**Table S17.** Evolutionary “fate” of the *Desmarestia herbacea* sex-determining genes in *Desmarestia dudresnayi*.

**Table S18.** Gene expression of the seven ancestral V-SDR genes in the reproductive stages of *Chordaria linearis* and *Desmarestia dudresnayi*.

**TableS19.** Expression of conserved VSDR genes (*Ectocarpus* sp. 7 SDR genes as reference) in *Fucus serratus* male and female mature receptacles.

**Table S20.** Transcriptional activity for V-SDR homologs in three Fucales species.

**Table S21.** Genomic data used in this study with metrics and accession numbers.

**Table S22.** Pairs of orthologous genes used to calculate the expected against observed number of orthologs per chromosome and to calculate rates of synonymous substitutions per site (*Ks*) for seven brown algal species.

**Table S23.** Gene age assignments based on phylostratigraphy as implemented in GenEra (https://github.com/josuebarrera/GenEra) for seven brown algal species, three plant species and one fungal species.

**Table S24.** Gene and repeat content for seven brown algal species over 100 kb sliding windows.

## SUPPLEMENTAL FIGURES

**Figure S1.**
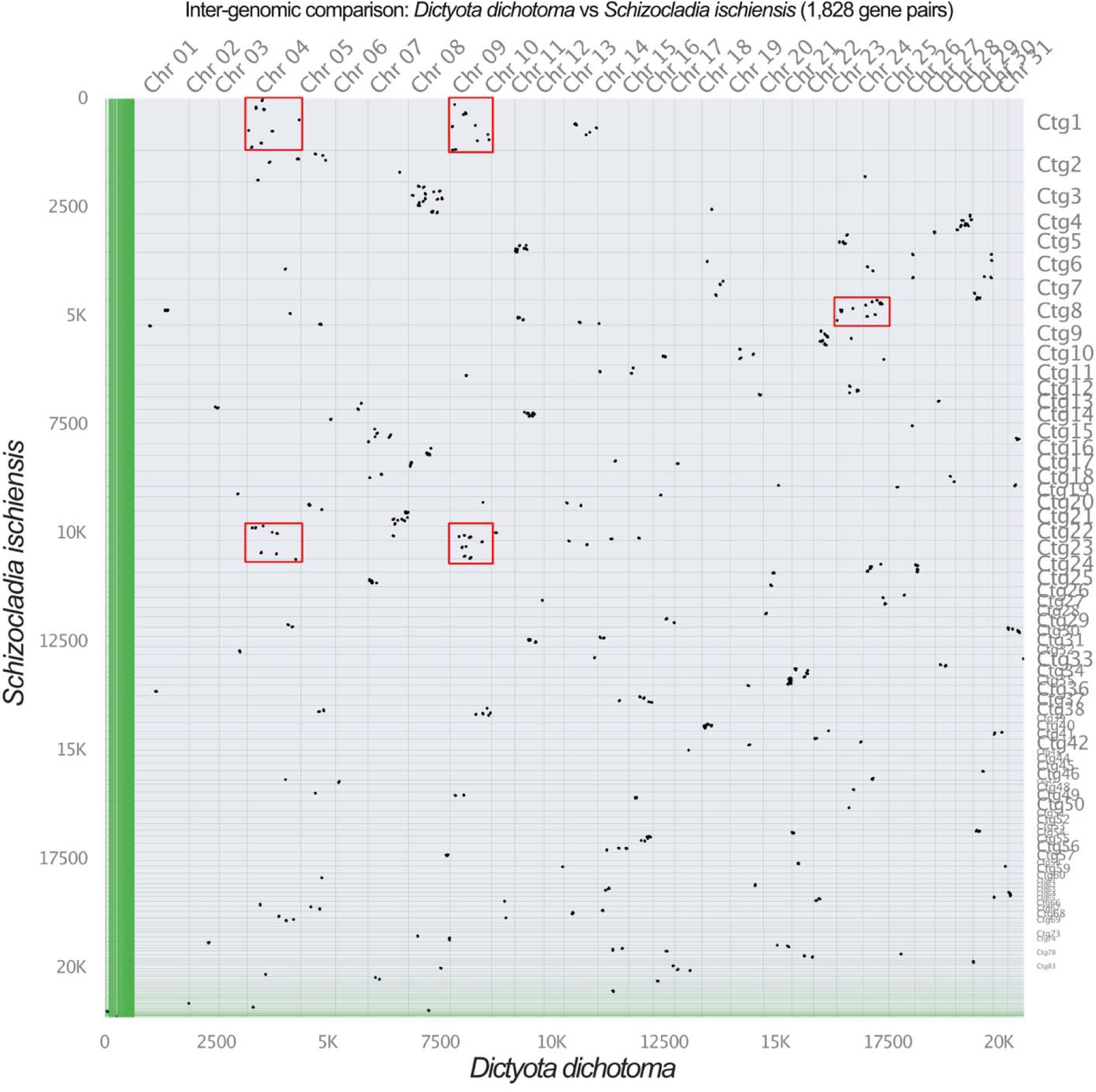
Macrosynteny plot between *S. ischiensis* and *D. dichotoma* using 1,828 orthologs. . We highlight two fusion-with-mixing events (red squares) between chromosomes 4 and 9, and between chromosomes 23 and 24 in *D. dichotoma*.

**Figure S2.**
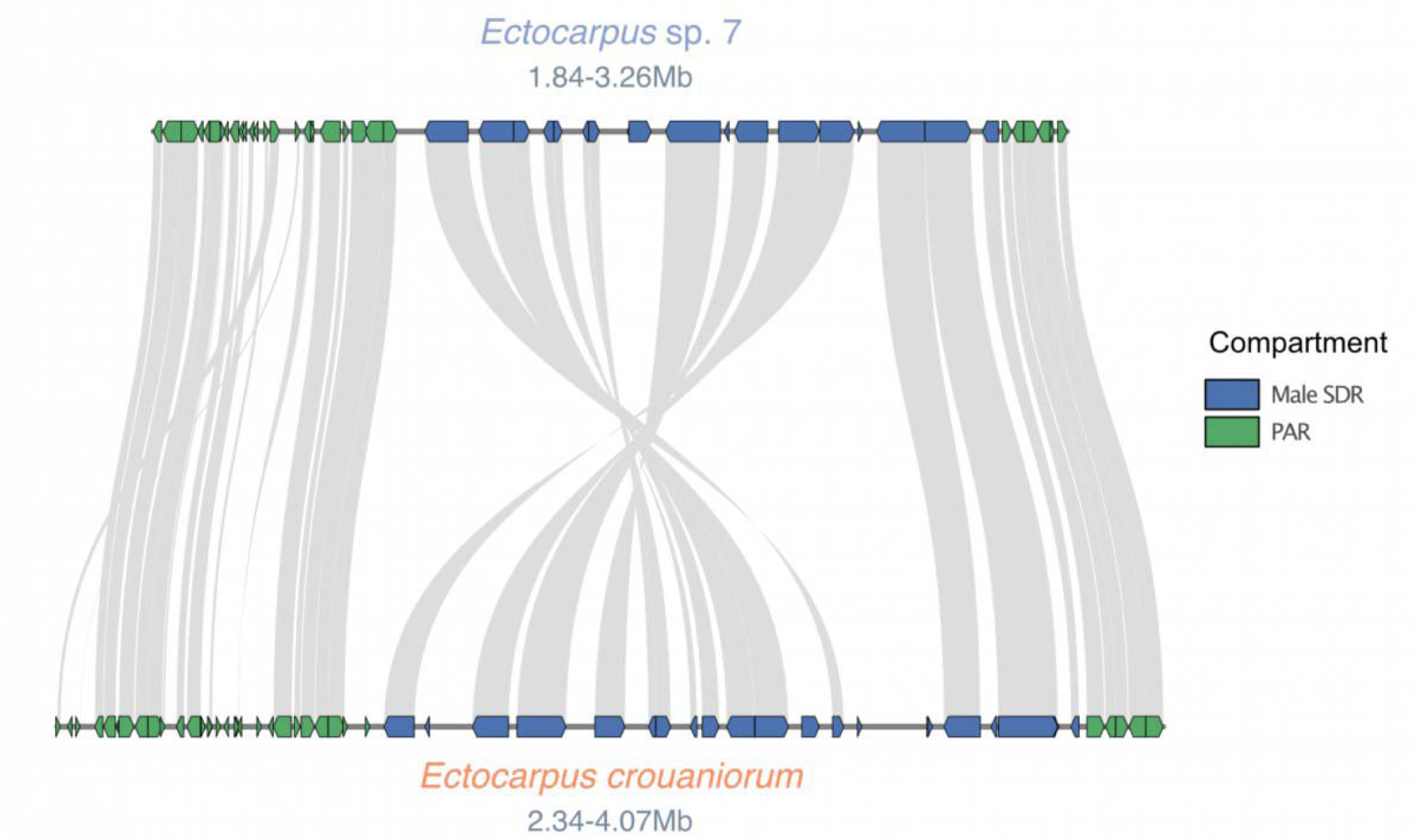
Male SDR synteny between *Ectocarpus* sp. 7 and *Ectocarpus crouaniorum*. One of the species underwent a recent inversion event within the SDR. The arrows in the boxes represent the orientation of each gene within the chromosome.

**Figure S3.**
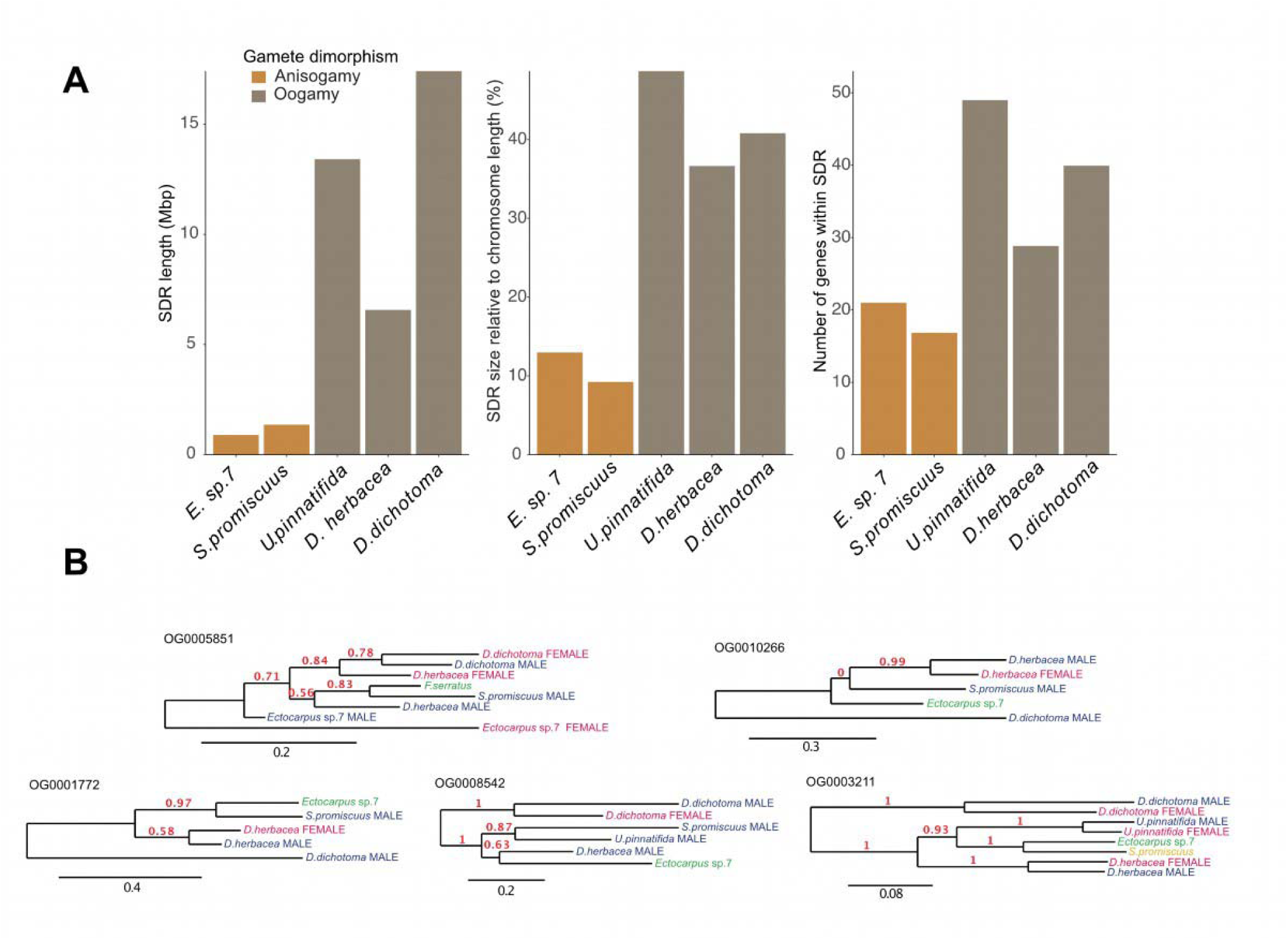
Detection of independently-acquired V-SDR genes across species. **(A)** Differences in the size of the male SDR between brown algal species based on the total sequence length, the relative size of the SDR compared to the length of the V chromosome and the number of protein-coding genes retained within the SDR. The bars are colored according to the level of gamete dimorphism in each species (based on ^24^). (B) Gene trees showing the independent acquisition of SDR gametologs across species that were previously interpreted as part of the ancestral male SDR genes.

**Figure S4.**
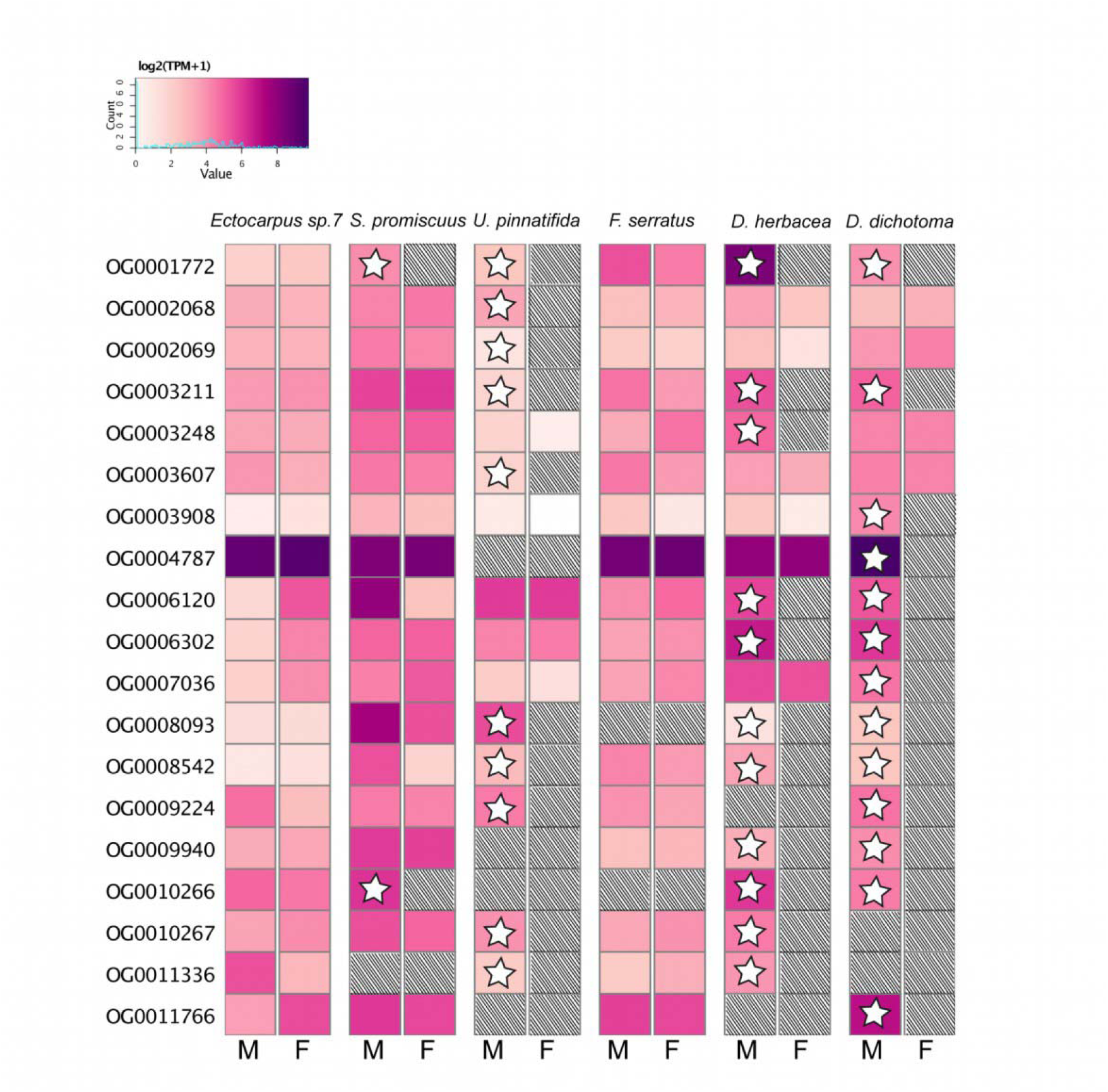
Expression of genes (log2(TPM+1)) that entered the SDR independently in different species. Expression is measured in mature male and female gametophytes, hashing marks missing orthologs, stars inside the cells indicate that the gene is inside the male non-recombining region (V-SDR). Orthogroups containing orthologs in less than three species or with multicopy genes were excluded from this analysis. M: male; F: female.

**Figure S5.**
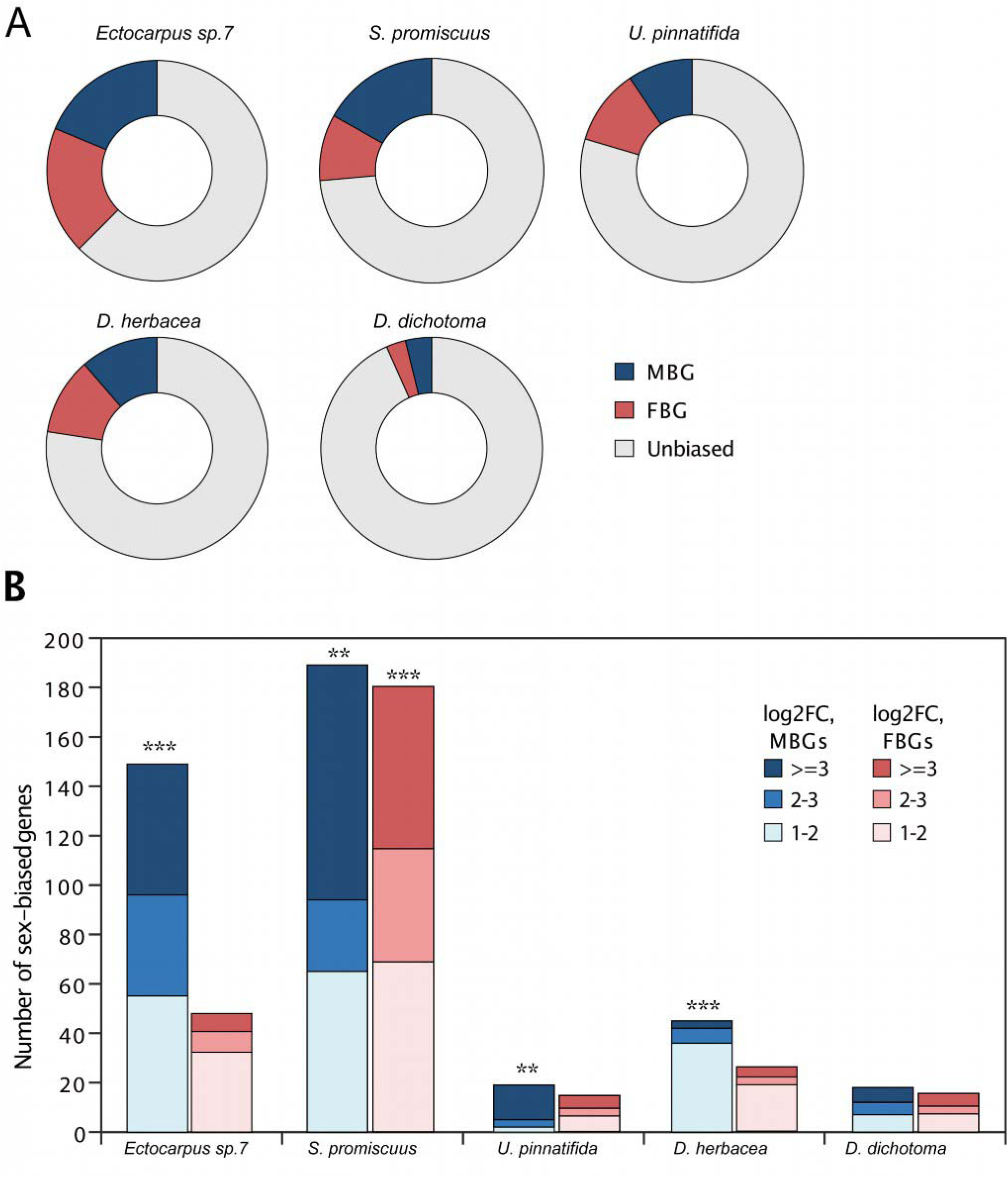
Sex-biased gene expression per dioicous species. (A) Proportion of sex biased genes in each of the five dioicous species. MBG: male-biased genes; FBG: female biased genes. (B) Number of sex-biased genes in the pseudoautosomal regions of sex chromosomes (U-V-SDRs excluded), male-biased genes are shown in blue and female-biased genes in red. Stars above the bars mark significant enrichment of the sex-biased genes on the PAR (Chi-square test, **p<0.01, ***p<0.001).

**Figure S6.**
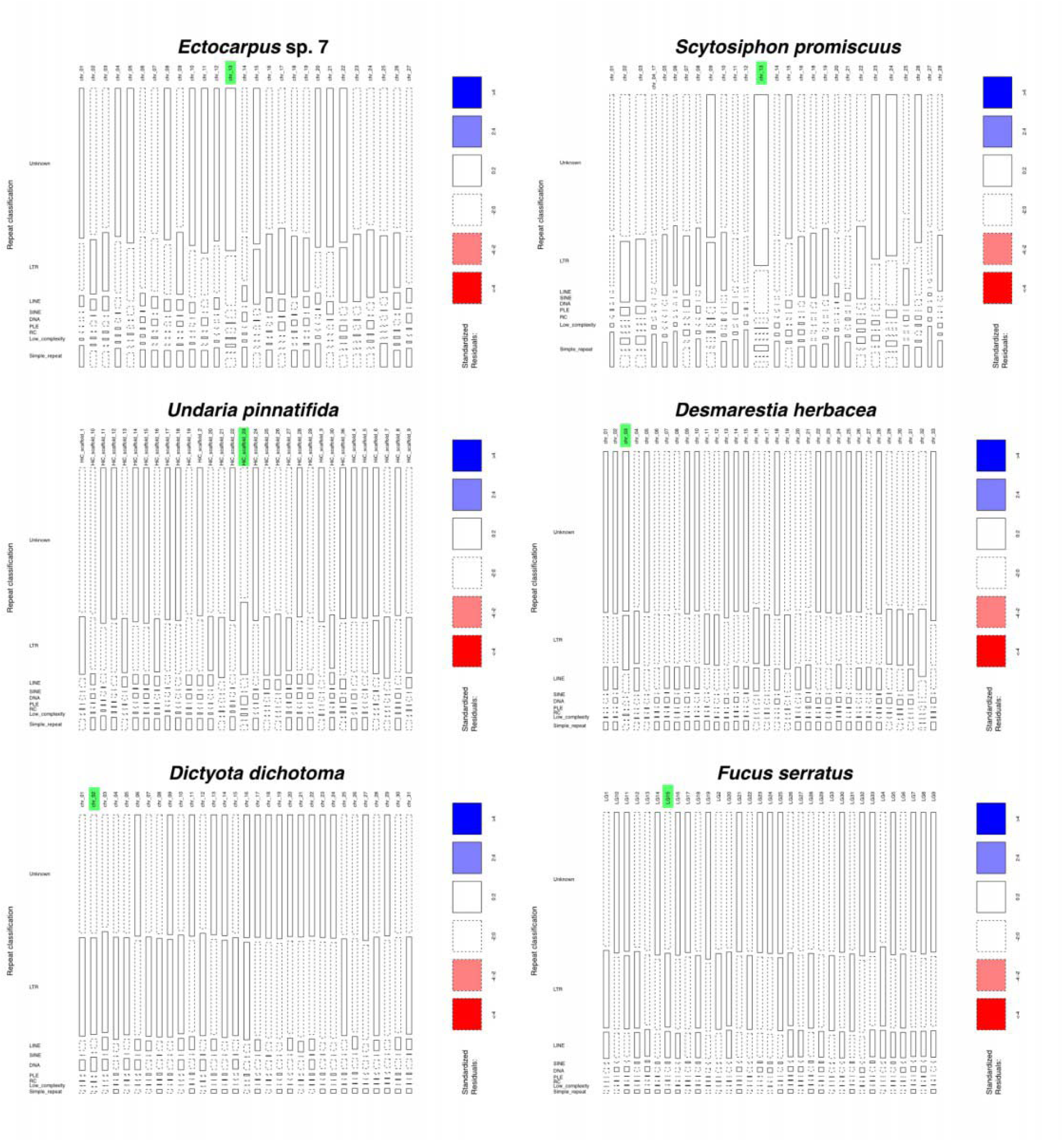
Distribution and proportion of classified transposable elements across the chromosomes of six brown algal species. The sex V chromosome or sex-homolog is highlighted in green. The residuals in the V sex chromosome were used to interpret the enrichment or depletion of Unclassified (Unknown) repetitive elements for each species.

**Figure S7.**
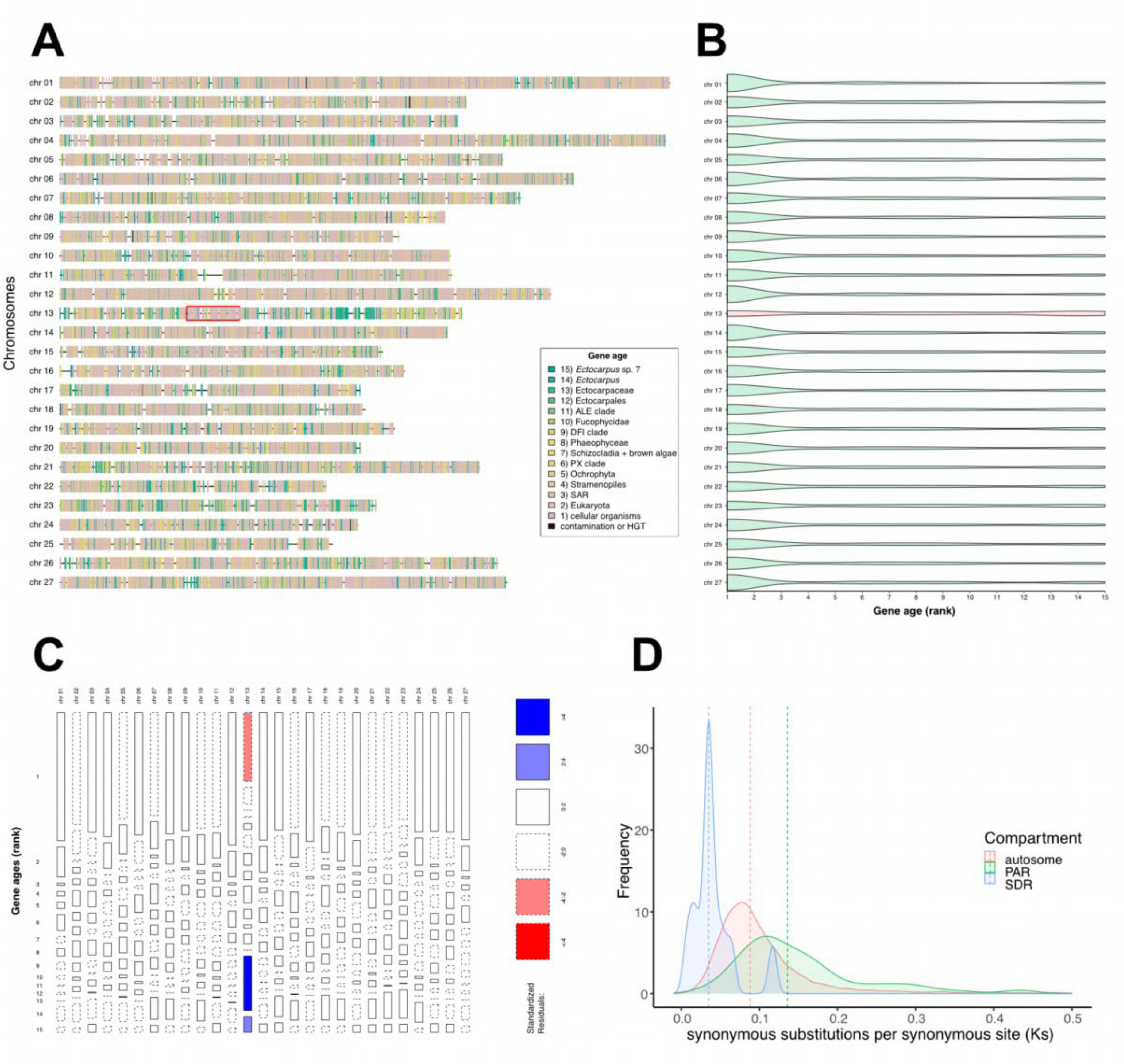
Gene ages across the *Ectocarpus* sp.7 genome. (A) Distribution of relative gene ages across the chromosomes of *Ectocarpus* sp. 7. The SDR of the V sex chromosome (chr 13) is highlighted with a red box. (B) The sex chromosome (red) has a significantly higher proportion of young genes and a lower proportion of old genes when compared to the autosomes (green; see Table S12). (C) Mosaic plot showing that the species-level (rank 15) and the genus-level (rank 14) genes are responsible for the enrichment of young genes in the sex chromosome. (D) The *Ks* values are significantly higher in the PARs of the sex chromosome when compared to the autosomes or the SDR (see Table S14).

**Figure S8.**
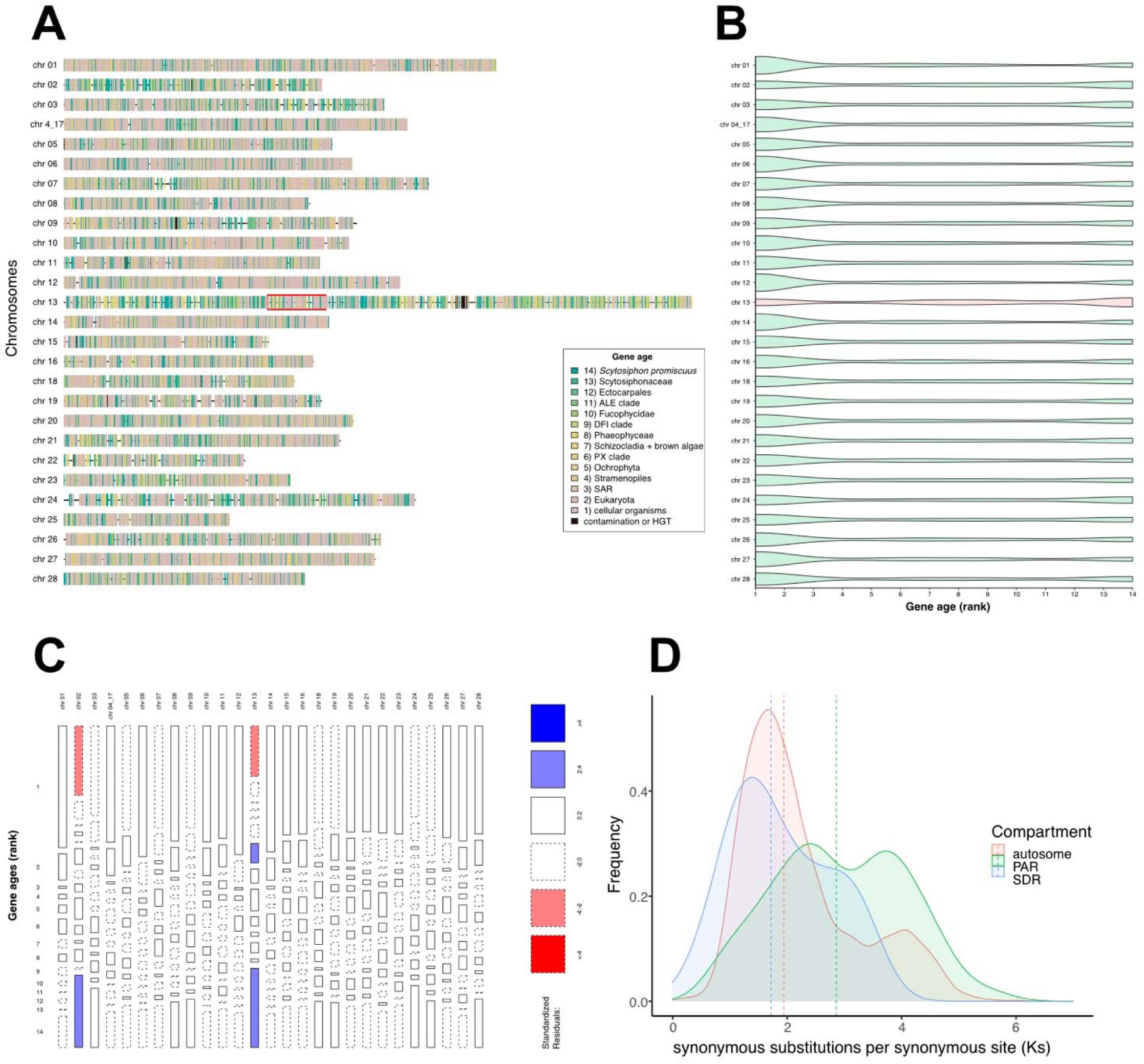
Gene ages across the *S. promiscuus* genome. A) Distribution of relative gene ages across the chromosomes of *Scytosiphon promiscuus*. The SDR of the V sex chromosome (chr 13) is highlighted with a red box. (B) The sex chromosome (red) has a significantly higher proportion of young genes and a lower proportion of old genes when compared to most of the autosomes (green; see Table S12). (C) Mosaic plot showing that the species-level (rank 14) genes are responsible for the enrichment of young genes in the sex chromosome. (D) The *Ks* values are significantly higher in the PARs of the sex chromosome when compared to the autosomes or the SDR (see Table S14).

**Figure S9.**
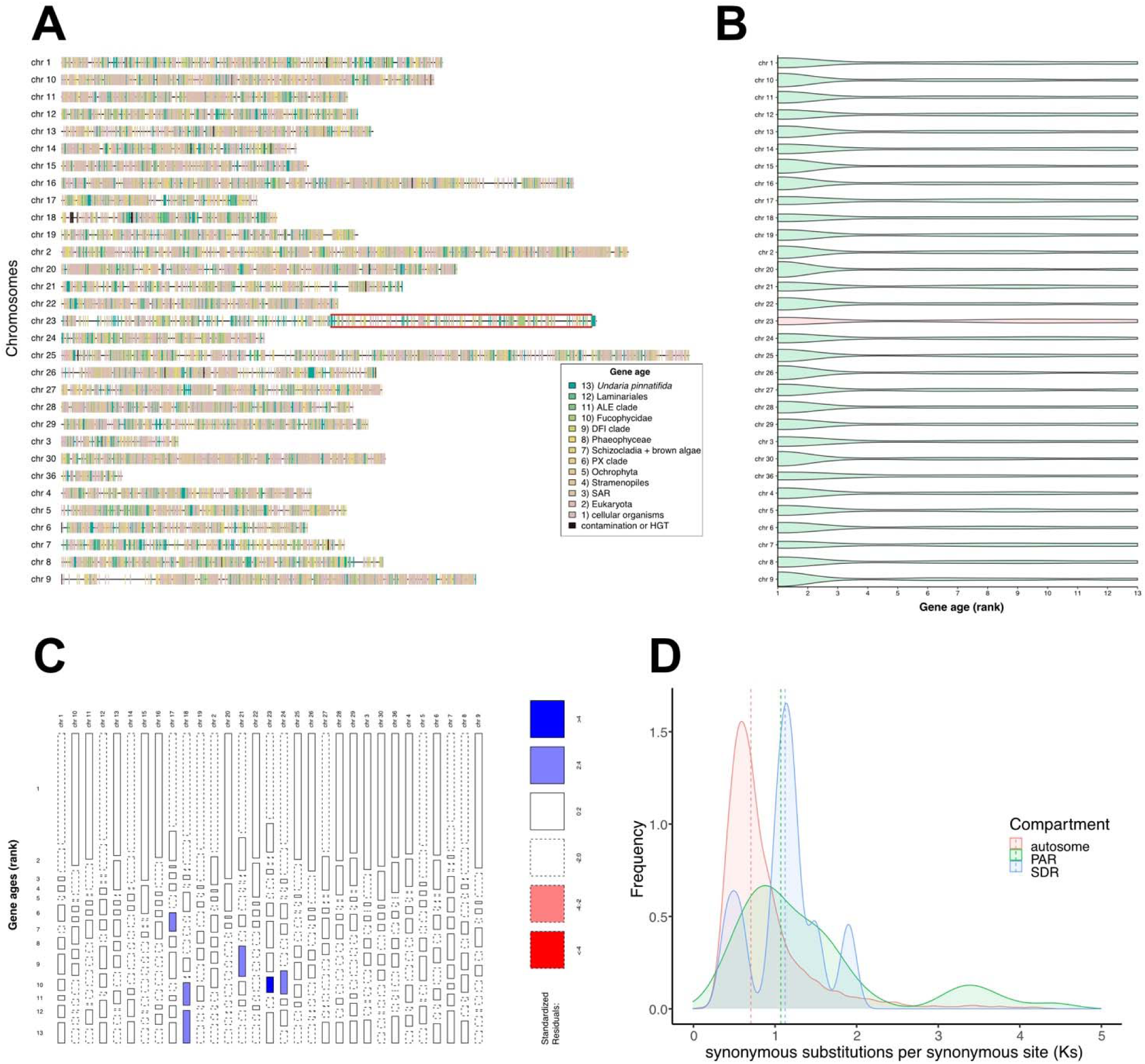
**Gene ages across the *U. pinnatifida* genome**. (A) Distribution of relative gene ages across the chromosomes of *Undaria pinnatifida*. The SDR of the V sex chromosome (chr 23) is highlighted with a red box. (B) The sex chromosome (red) has a significantly higher proportion of young genes and a lower proportion of old genes when compared to most of the autosomes (green; see Table S12). (C) Mosaic plot showing that the ALE-clade genes (rank 11) are responsible for the enrichment of young genes in the sex chromosome. (D) The *Ks* values are significantly higher in the sex chromosome when compared to the autosomes (see Table S14), showing similar values in the PARs and in the SDR.

**Figure S10.**
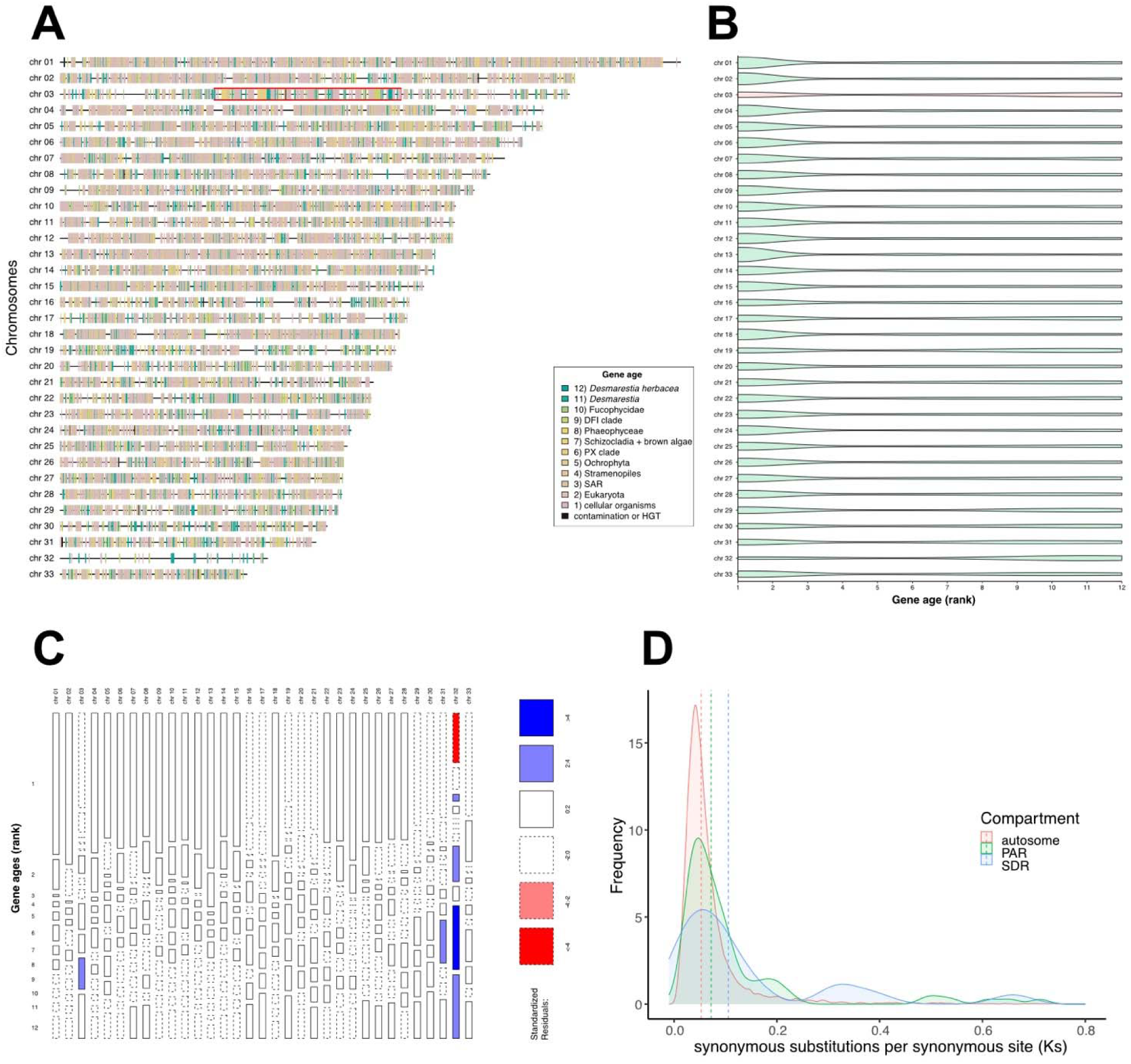
Gene ages across the *D. herbacea* genome. . (A) Distribution of relative gene ages across the chromosomes of *Desmarestia herbacea*. The SDR of the V sex chromosome (chr 03) is highlighted with a red box. (B) The sex chromosome (red) has a significantly higher proportion of young genes and a lower proportion of old genes when compared to most of the autosomes (green; see Table S12). (C) Mosaic plot showing that the genus-level (rank 11) genes are responsible for the enrichment of young genes in the sex chromosome. (D) The *Ks* values are significantly higher in the sex chromosome when compared to half of the autosomes (see Table S14). Non-significance of *Ks* values across chromosomes may be driven by the conflation with the *Ks* values in *Desmarestia dudresnayi*. The SDR displays higher *Ks* values compared to the PARs or the autosomes.

**Figure S11.**
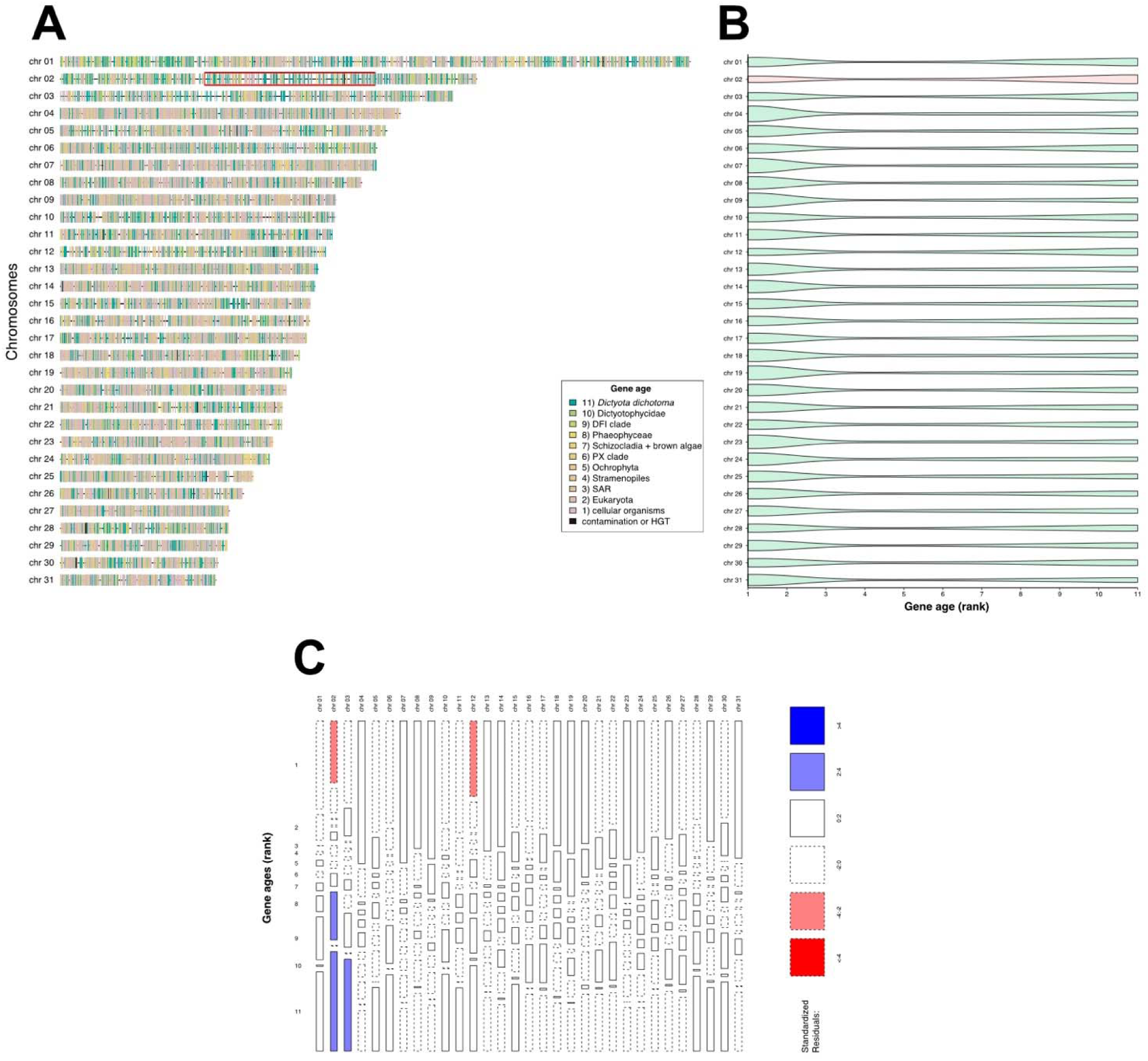
Gene ages across the *D. dichotoma* genome. (A) Distribution of relative gene ages across the chromosomes of *Dictyota dichotoma*. The SDR of the V sex chromosome (chr 02) is highlighted with a red box. (B) The sex chromosome (red) has a significantly higher proportion of young genes and a lower proportion of old genes when compared to most of the autosomes (green; see Table S12). (C) Mosaic plot showing that the species-level (rank 11) and the DFI-clade-level (rank 9) genes are responsible for the enrichment of young genes in the sex chromosome. *Ks* values were not calculated for *D. dichotoma*, due to a saturation of synonymous mutations with the closest species in the PhaeoExplorer database (*Halopteris paniculata*).

**Figure S12.**
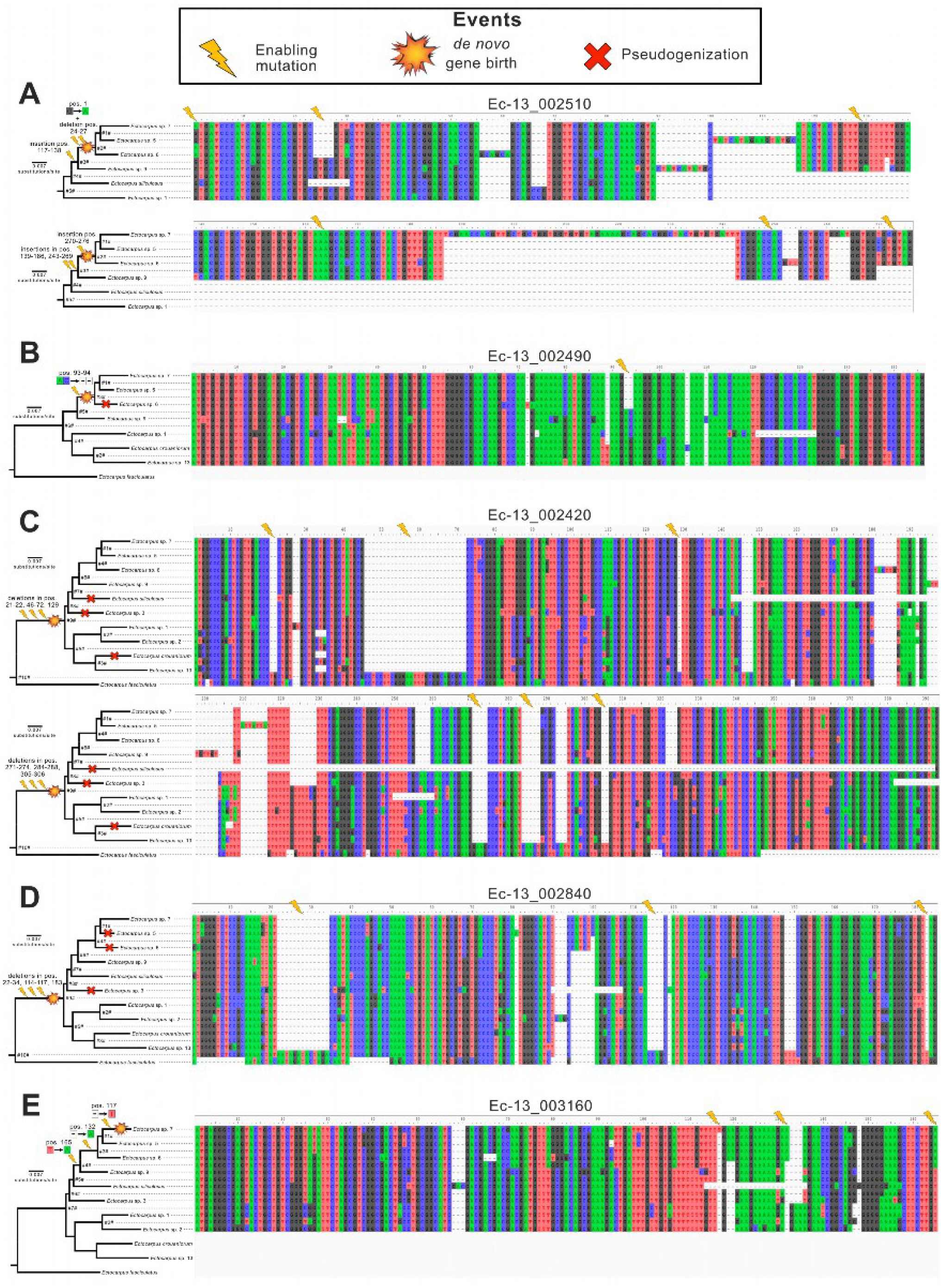
Ancestral state reconstruction behind five *de novo* gene birth events in *Ectocarpus*. (A) Evolutionary steps leading to the emergence of gene Ec-13_002510. The first step is a big insertion (pos. 117-269) in the last common ancestor between *E.* sp. 7 and *E.* sp. 9 (node #3#). Later, on the common ancestor between *E*. sp. 7 and *E*. sp 6 (node #2#), a G-to-A substitution led to the emergence of a start codon, an insertion on the 3’ end (pos. 270-276) introduced a stop codon and four-nucleotide deletion (pos. 24-27) led to a triadic sequence (i.e., a multiple-of-three nucleotide sequence). (B) Evolutionary steps leading to the emergence of gene Ec-13_002490. A two-nucleotide deletion (pos. 93-94) led to a triadic sequence with an open reading frame (ORF) in the common ancestor of *E*. sp. 7 and *E*. sp. 6 (node #3#). A later insertion (pos. 72) led to the disruption of this gene in *E*. sp. 6. (C) Evolutionary steps leading to the emergence of gene Ec-13_002420. The last common ancestor between *E*. sp. 7 and *E. crouaniorum* (node #9#) experienced six deletions (pos. 21-22, 46-72, 129, 271-274, 284-288 and 305-306) that led to the emergence of a triadic sequence with an ORF. Later deletions led to the independent disruption of this gene in *E. siliculosus* (pos. 143-392), *E*. sp. 3 (pos. 382-393) and *E. crouaniorum* (pos. 206-207). (D) Evolutionary steps leading to the emergence of gene Ec-13_002840. The last common ancestor between *E*. sp. 7 and *E. crouaniorum* (node #9#) experienced three deletions (pos. 22-34, 114-117 and 183) that led to the emergence of a triadic sequence with an ORF. Later mutations led to the independent disruption of this gene in *E*. sp. 5 (pos. 81), *E*. sp. 6 (pos. 1-2) and *E*. sp. 3 (pos. 90-107). (E) Evolutionary steps leading to the emergence of gene Ec-13_003160. The common ancestor of *E*. sp. 7 and *E*. sp. 9 (node #4#) experienced a T-to-A substitution (pos. 165) that led to the emergence of a stop codon. The common ancestor between e. sp. 7 and E. sp. 6 (node #3#) later experience a deletion (pos. 132) that shifted the sequence towards a triadic pattern. Finally, an additional insertion (117) established the triadic pattern within the sequence, leading to the emergence of an ORF in *E*. sp. 7.

**Figure S13.**
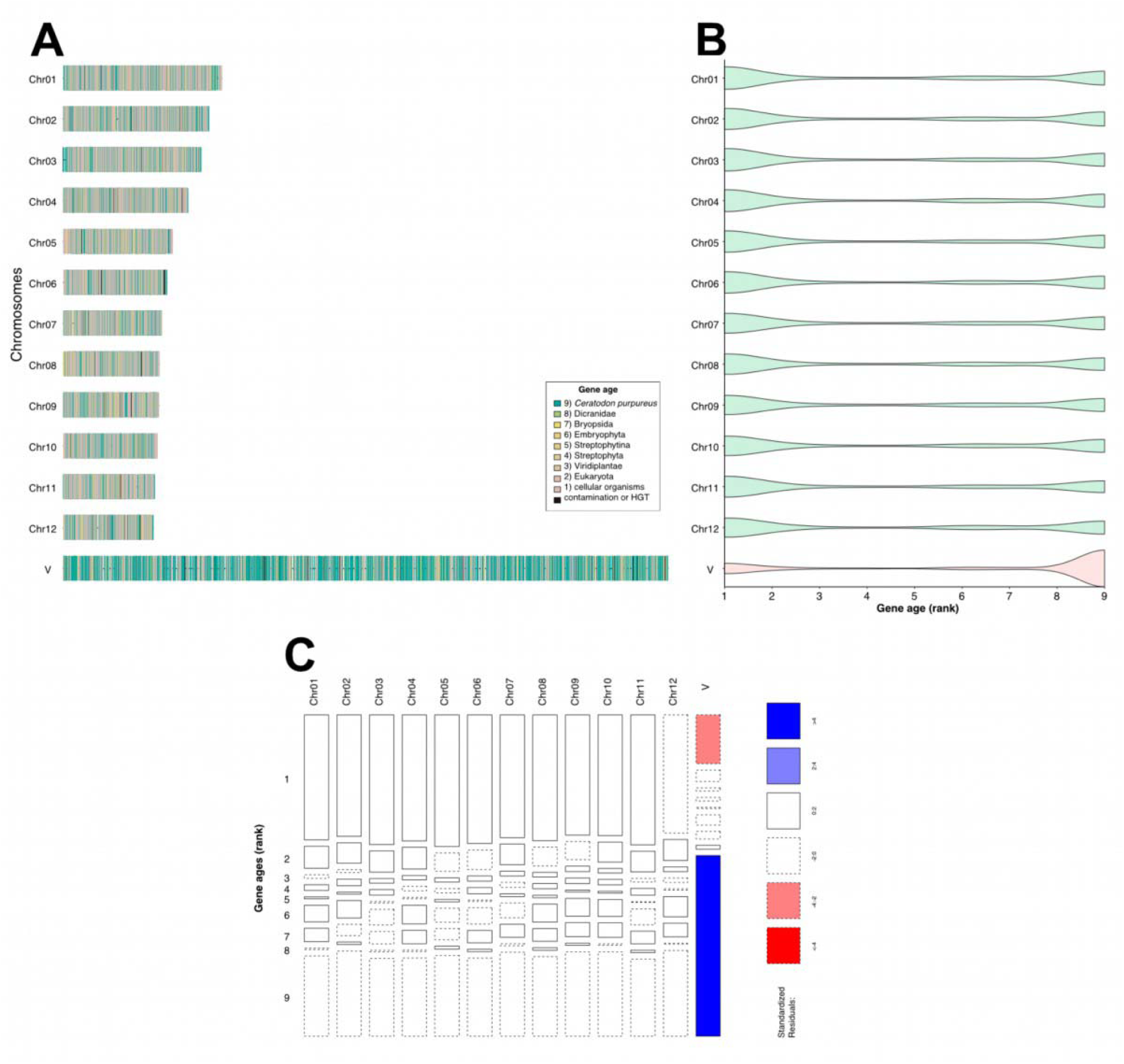
Gene ages across the *Ceratodon purpureum* genome. (A) Distribution of relative gene ages across the chromosomes of *Ceratodon purpureum*. (B) The V sex chromosome (V; red) has a significantly higher proportion of young genes and a lower proportion of old genes when compared to the autosomes (green; see Table S12). (C) Mosaic plot showing that the species-level genes (rank 9) are responsible for the enrichment of young genes in the sex chromosome.

**Figure S14.**
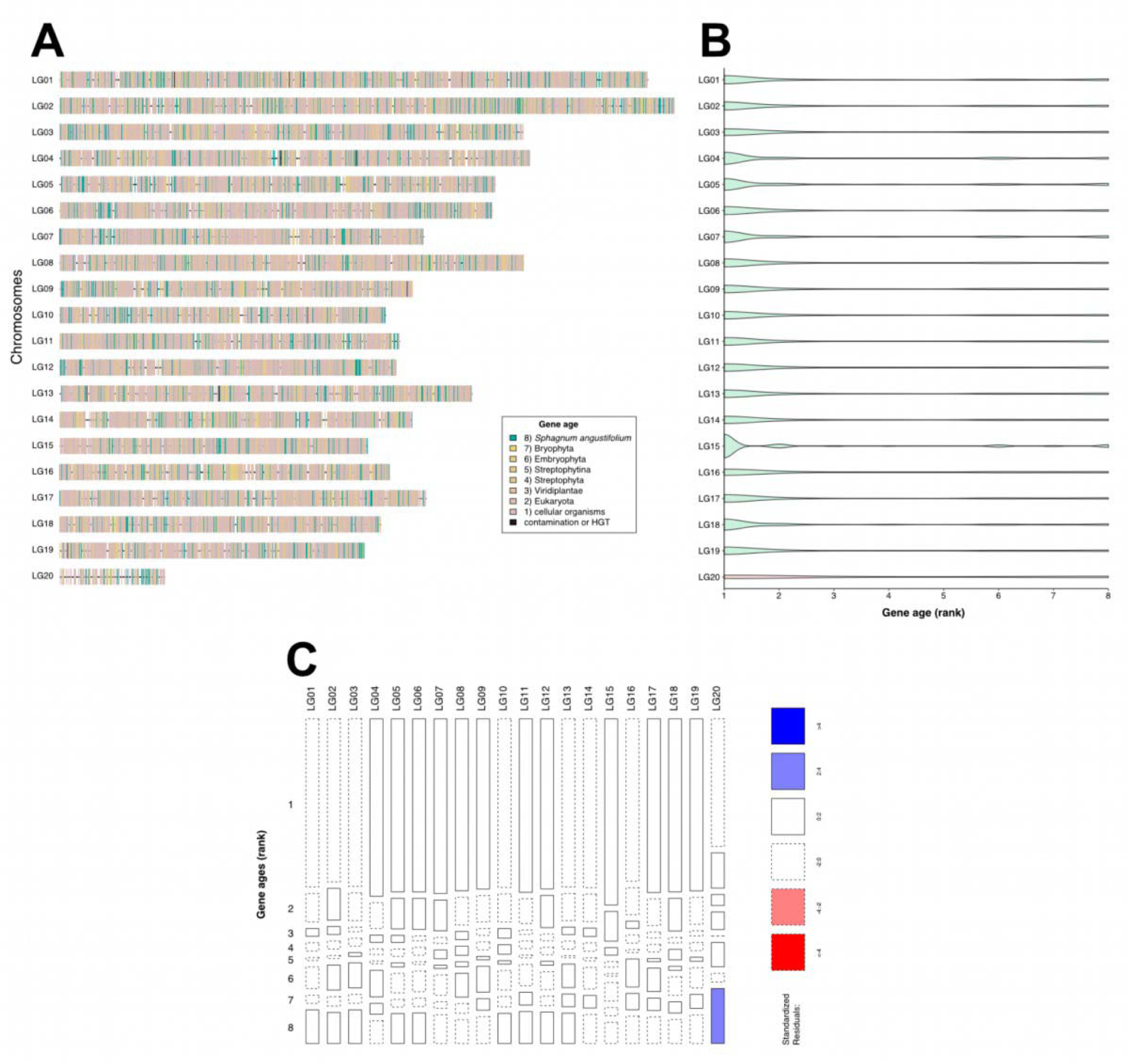
Gene ages across the *Sphagnum angustifolium* genome. (A) Distribution of relative gene ages across the chromosomes of *Sphagnum angustifolium*. (B) The V sex chromosome (LG20; red) has a significantly higher proportion of young genes and a lower proportion of old genes when compared to the autosomes (green; see Table S12). (C) Mosaic plot showing that the species-level genes (rank 8) are responsible for the enrichment of young genes in the sex chromosome.

**Figure S15.**
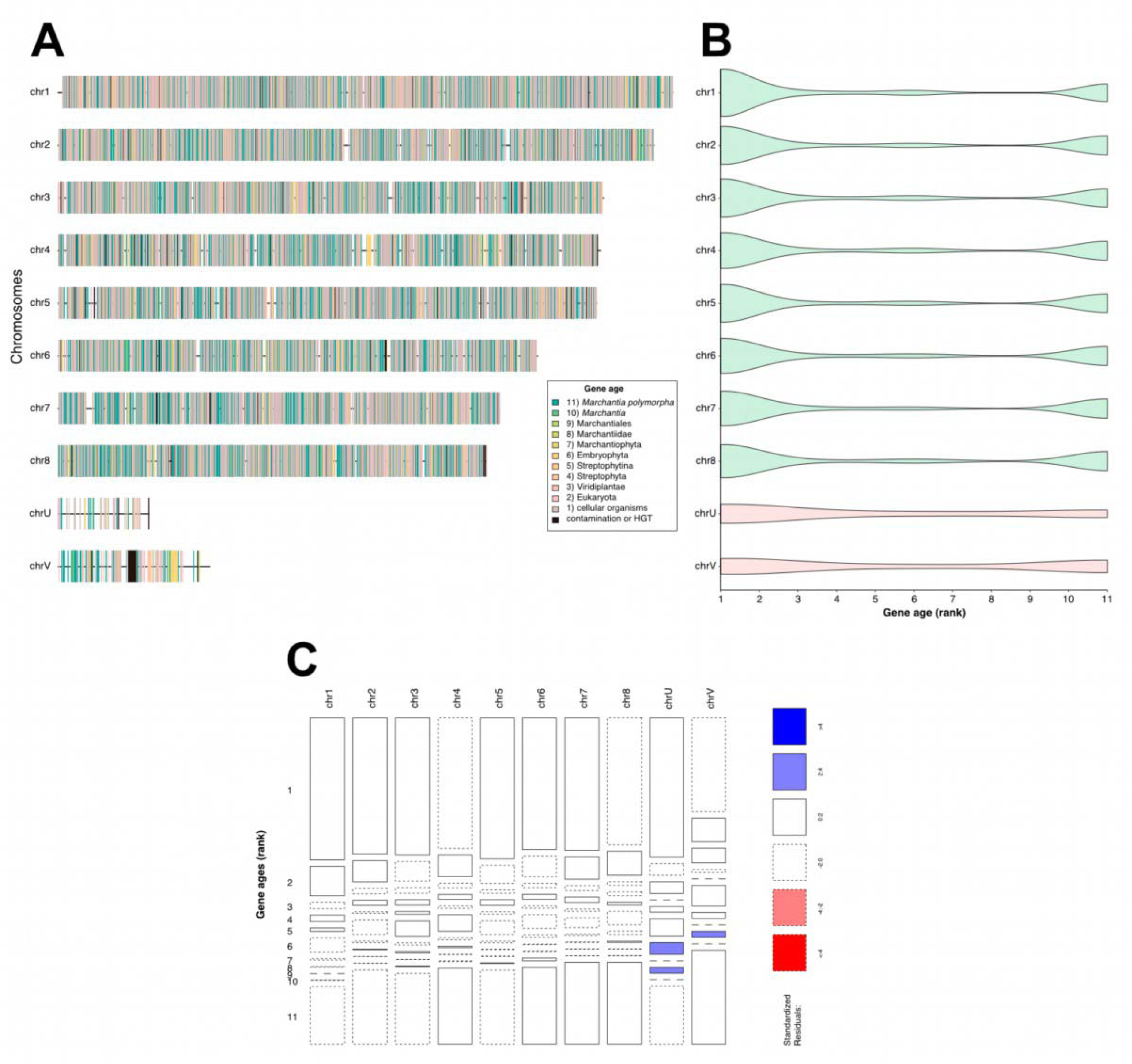
Gene ages across the *Marchantia polymorpha* genome. (A) Distribution of relative gene ages across the chromosomes of *Marchantia polymorpha*. (B) The U/V sex chromosomes (chrU and chrV; red) show non-significant differences in gene age distribution when compared to the rest of the chromosomes (green; see Table S12). (C) Mosaic plot showing non-significant differences between the sex chromosomes and the autosomes.

**Figure S16.**
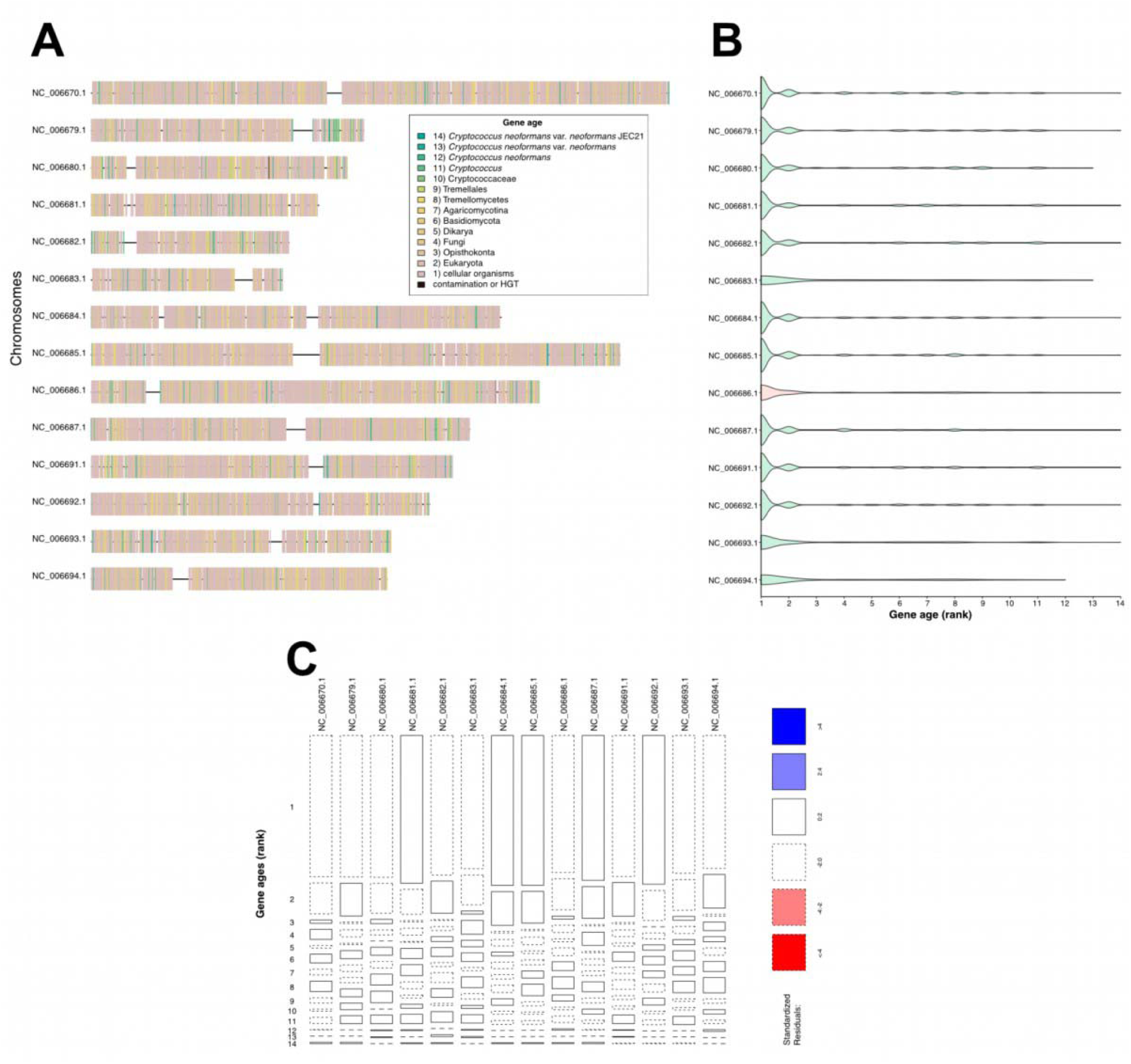
Gene ages across the *Cryptoccocus neoformans* var. *neoformans* JEC21 genome. (A) Distribution of relative gene ages across the chromosomes of *Cryptoccocus neoformans*. (B) The mating-type chromosome (NC_006686.1; red) shows non-significant differences in gene age distribution when compared to the rest of the chromosomes (green; see Table S12). (C) Mosaic plot showing no discernible pattern of gene age distribution in any of the chromosomes.

**Figure S17.**
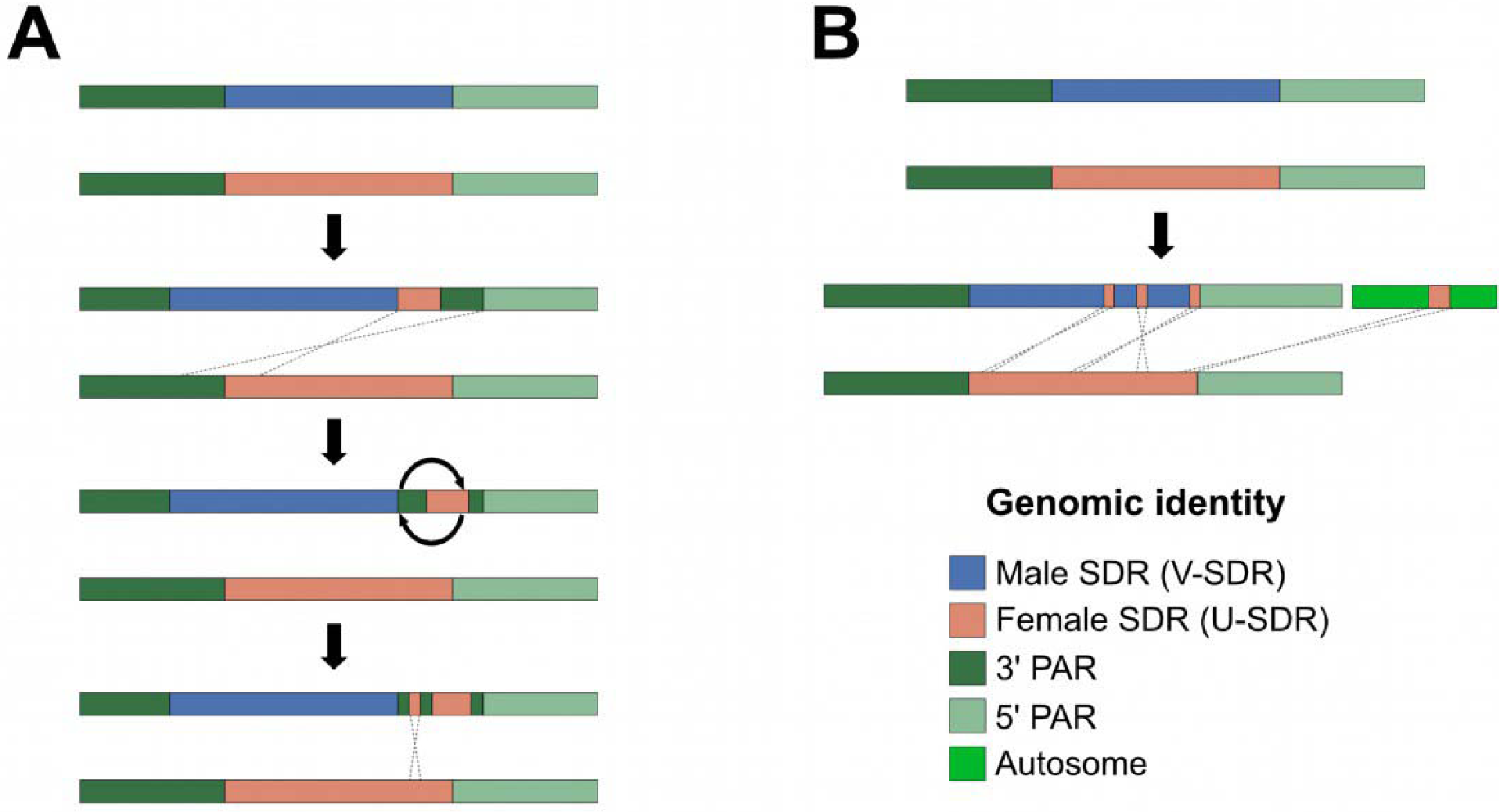
Proposed scenarios for the transition from dioicy to monoicy in *Chordaria linearis* and *Desmarestia dudresnayi*. (A) The ancestor of *Chordaria linearis* likely underwent an initial translocation event from the U chromosome to the V chromosome, inserting part of the U-SDR and a piece of the 3’ PAR towards the 5’ end of the V-SDR through a non-homologous recombination event. A subsequent inversion within this translocation spread the 3’PAR genes to both sides of the U-SDR insertion. Finally, a second non-homologous recombination event inserted an additional piece of the U-SDR within the 3’ PAR translocation. (B) The ancestor of *Desmarestia dudresnayi* underwent three translocations of U-SDR genes into the V-SDR. Additionally, a fourth translocation event happened between the U-SDR and an autosome (chr_04).

**Figure S18.**
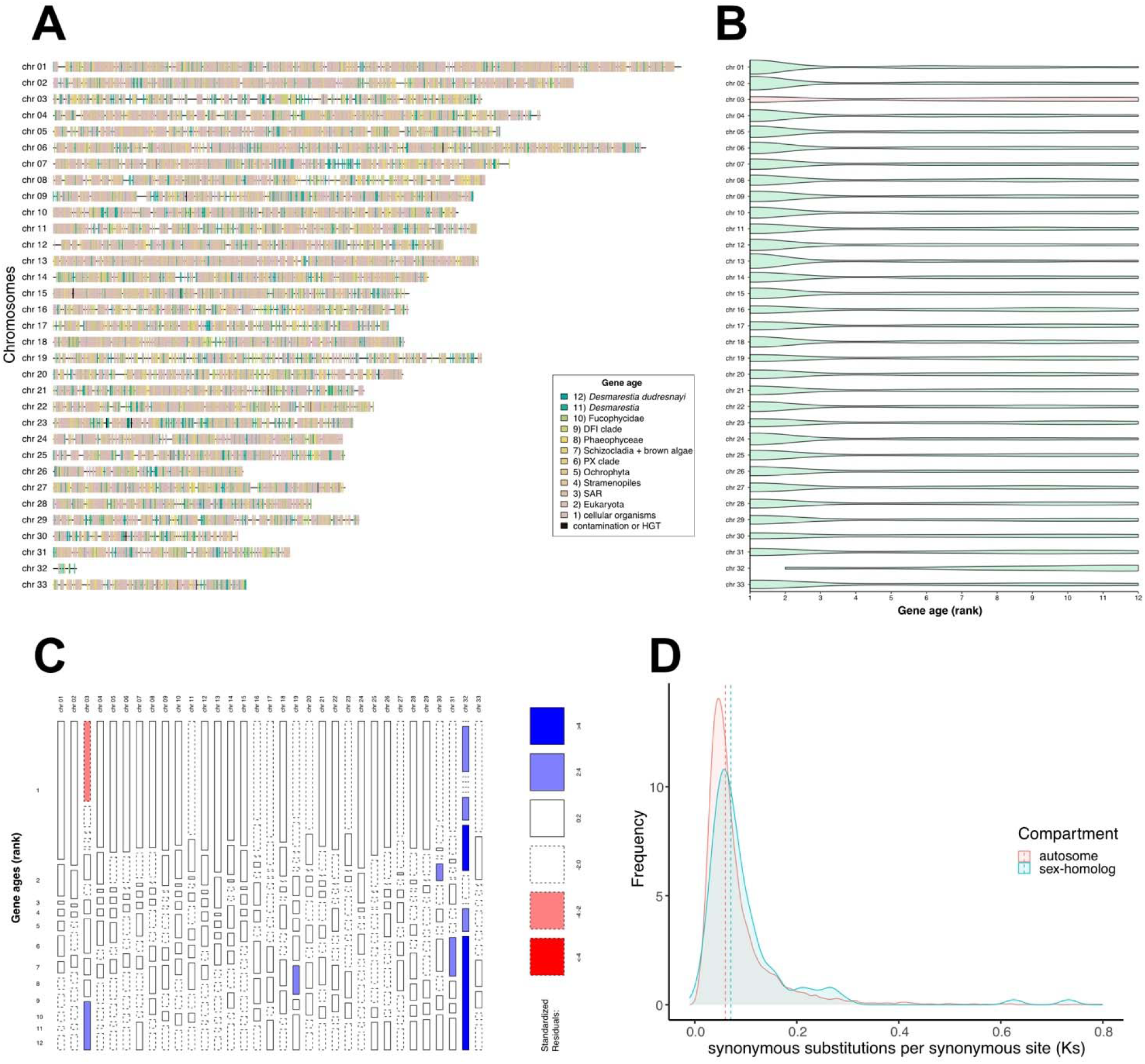
Gene ages across the *D. dudresnayi* genome. . (A) Distribution of relative gene ages across the chromosomes of *Desmarestia dudresnayi*. (B) The sex-homolog in *D. dudresnayi* (chr 03; red) has a significantly higher proportion of young genes and a lower proportion of old genes when compared to most of the other chromosomes (green; see **Table S12**). (C) Mosaic plot showing that the species-level genes (rank 12) are responsible for the enrichment of young genes in the sex-homolog. (D) The *Ks* values are similar in the sex-homolog when compared to the other chromosomes (see **Table S14**).

**Figure S19.**
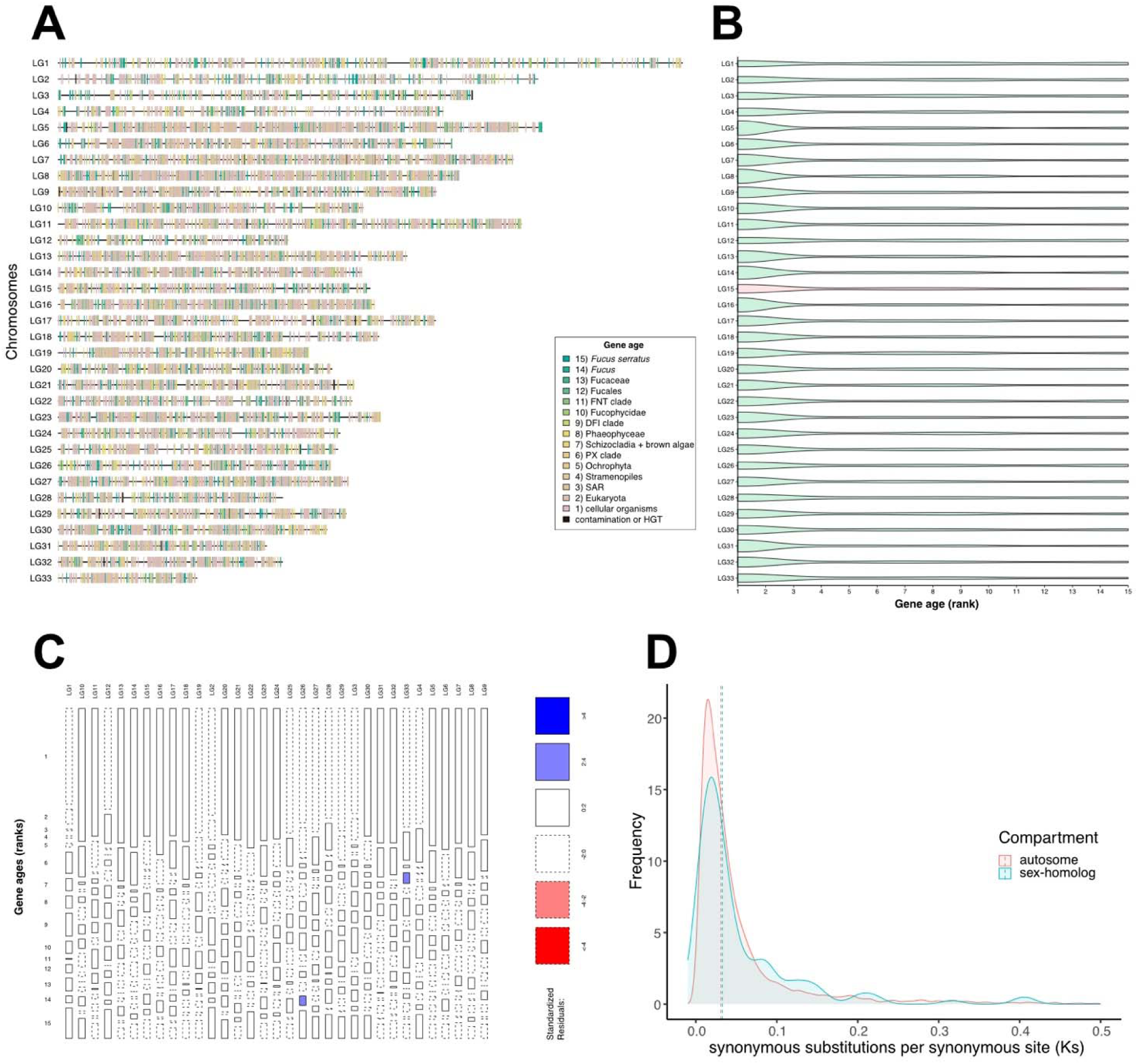
Gene ages across the *F. serratus* genome. . (A) Distribution of relative gene ages across the chromosomes of *Fucus serratus*. (B) The sex-homolog in *F. serratus* (LG15; red) shows no significant differences in gene age distribution when compared to the rest of the chromosomes (green; see **Table S12**). (C) Mosaic plot showing no discernible pattern of gene age distribution in any of the chromosomes. (D) The *Ks* values are similar in the sex-homolog when compared to the other chromosomes (see **Table S14**).

## METHODS

### Resource availability

#### Lead contact

Further information and requests for resources and reagents should be directed to and will be fulfilled by the lead contact, Susana M. Coelho.

#### Data and code availability

- The accession numbers and download links for all the genomic data that was generated and used in this study are available on **Table S21**.
- This paper does not report original code.

#### Biological material

*Scytosiphon promiscuus, Dictyota dichotoma*, *Undaria pinnatifida* and *Desmarestia dudresnayi* haploid gametophytes were cultivated in the laboratory conditions as in ^83^. We cultivated the gametophytes at 14°C with a photoperiod of 12:12 h light:dark an irradiance of 25µmol photons.m-2.s-1. The media consisted of filtered natural seawater (NSW), which was autoclaved and enriched with half-strength Provasoli nutrient solution (Provasoli-enriched seawater; PES)^83^. We grew the first biomass in 140mm Petri dishes and the gametophytes were later transferred to 1L flask with gentle aeration. The gametophytes were fragmented once a month and the media were changed every two weeks to promote biomass production. Prior to freezing, gametophytes were treated with antibiotics for 3 days with a gentle agitation and under the same culture conditions. The first day, gametophytes were treated with a mix Streptomycin (2g/L of PES), Penicillin G (0.5g/L of PES) and Chloramphenicol (0.1g/L of PES); the next day with Ampicilin (1g/L of PES) and finally the last day with Kanamycin (1g/L of PES). Between each day of treatment and before freezing, gametophytes were rinsed with 500mL of NSW to remove the traces of antibiotic.

Samples for fucoid algae sexual and vegetative tissue were collected in the intertidal zone during low tides in June 2012 from Viana do Castelo (*F. vesiculosus*, *A. nodosum*) and Caminha (Rio Minho; *F. ceranoides*), northern Portugal. Sexual phenotypes were verified in the field by sectioning and observing receptacles under a field microscope. Tissue samples were flash-frozen in liquid nitrogen on the shore and transported to the laboratory in a cryoshipper, after which they were lyophilized and stored dry at room temperature on silica crystals. See **Table S21** for list of strains used in this study.

#### DNA and RNA extraction and sequencing

Genomic DNA was isolated from algal tissue (∼100mg) by grinding into fine powder under liquid nitrogen and subsequent cell lysis in 500μL of Genomic Lysis Buffer (OMNIPREP for plant kit) for 1 hour at 60°C. The lysate was cleaned up with 200μL of chloroform and DNA was precipitated in EtOH. The DNA pellet was digested in CF buffer (Macherey-Nagel) for 45 min at 65°C and purified using NucleoBond AXG20 Mini columns according to the user manual (Macherey-Nagel). Final high molecular weight gDNA was quantified (Qubit), analyzed for purity (Nanodrop) and checked for size distribution (Femto Pulse System) before preparing the sequencing libraries. We sequenced the libraries using an Oxford Nanopore Technologies (ONT) MinION Mk1B. We prepared the ONT libraries using an SQK-LSK110 library preparation kit for R9.4.1 flow cells and an SQK-LSK114 library preparation kit for R10.4.1 flow cells. Two libraries were sequenced for *Desmarestia dudresnayi* on R9.4.1 flowcells and a third library was sequenced on a R10.4.1 flowcell.

RNA was isolated from mature gametophytes of *Undaria pinnatifida* and *Scytosiphon promiscuus* following modified procedure of Qiagen RNAeasy kit and the TruSeq RNA Library Prep Kit v2 was used to sequence the transcriptomes in an Illumina NextSeq 2000 platform (150bp, PE reads). Extraction of total RNA from fucoid algae (*F. vesiculosus, A. nodosum* and *F. ceranoides*) was performed following 65 and RNA libraries were sequenced on Illumina HiSeq 2000 machine (100 bp, PE reads).

#### Genome assembly and annotation

Whole-genome assemblies and annotations of *S. promiscuus* male, *D. dichotoma* male, *D. herbacea* male and female, *E. crouanorium* male, *C. linearis*, *S. ischiensis* and *F. serratus* male were obtained from Denoeud et al.^26^. We also downloaded the genome of *Ectocarpus* sp. 7^27^ and the male genome of *Undaria pinnatifida*^84^ which were already assembled at a chromosome level. For *Desmarestia dudresnayi*, we performed genome sequencing, *de novo* genome assembly and *ab initio* gene annotation. Base calling was done using ONT Guppy^85^ with the configuration files dna_r9.4.1_450bps_sup.cfg and dna_r10.4.1_e8.2_400bps_sup.cfg and the options --trim_adapters –trim_primers, yielding 17.4 Gbp of data in 2,871,152 reads. We merged all the reads and analyzed them using Kraken v2.1.2^86^ and the bacteria database (downloaded 08-2022) to remove potential contaminant sequences. All data classified as bacterial reads by Kraken were screened using blastN v2.13.0+^87^ (-evalue 0.001 -num_alignments 20) against the NCBI genbank bacterial database (downloaded 11-2023). The blastN output was visualized in MEGAN v6.23.4^88^, and all the reads that were declared as bacterial were extracted and removed from further analyses. We obtained 1,908,772 decontaminated reads with an average length of 5.1Kbp (9.8 Gbp of data, 20x coverage), which were deposited on the NCBI Sequence Read Archive (see **Table S21**).

The decontaminated reads were assembled *de novo* using flye v2.9.1-b1780^89^ with the options ‘--nano-raw - g 450m -t 28 -i 3 --scaffold’. The draft assembly consisted of 1,032 contigs with a total size of 425 Mbp, an N50 of 4.6 Mbp and an L50 of 29 contigs. We used TransposonPSI (http://transposonpsi.sourceforge.net/) to predict the transposable elements and RepeatScout v1.0.6^90^ to predict the simple repeats in the genome assembly. Both predictions were combined to soft-mask the repetitive content in the genome assembly using bedtools maskfasta v2.27.1^91^. We mapped the RNA-seq data of *Desmarestia dudresnayi* from the PhaeoExplorer database^26^ to the soft-masked genome assembly using STAR v2.7.1a^92^. We used BRAKER v2.1.6 alongside the RNA-seq data^93^ to predict the protein-coding genes in the soft-masked genome assembly.

#### Hi-C library preparation and sequencing for chromosome-level assemblies

We generated Hi-C libraries for three male genomes (*Scytosiphon promiscuus*, *Desmarestia herbacea* and *Dictyota dichotoma*) and two female genomes (*Ectocarpus* sp. 7 and *Desmarestia herbacea*). Fresh algal tissue was cross-linked for 20 minutes at room temperature in a solution of 2% formaldehyde with filtered natural sea water (NSW) and then transferred into a 400 mM Glycine solution with filtered NSW for five minutes to quench the formaldehyde. The samples were then stored at −80°C until use. The Hi-C libraries were prepared as follows. The samples were de-frosted in 1 mL of 1x *Dpn*II buffer with protease inhibitors (Roche cOmplete™), transferred to Precellys VK05 lysis tubes (Bertin Technologies, Rockville, MD) and disrupted using the Precellys apparatus with five grinding cycles of 30 seconds at 7,800 rpm followed by 20 second pauses. SDS was added to the lysate at 0.5% final concentration and samples were incubated for 10 minutes at 62°C, followed by the addition of Triton-X100 to a final concentration of 1% and 10 minutes of incubation at 37°C under gentle shaking. We added 500 U of *Dpn*II to 4.6 mL of the digestion mixture and incubated the samples for two hours at 37°C under gentle shaking (180 rpm in an inclined rack to prevent sedimentation), followed by the addition of another 500 U of *Dpn*II and an overnight incubation under the same conditions. The digested samples were centrifuged at 4°C for 20 minutes at 16,000×g. The supernatant was discarded and the pellet was incubated for biotinylation at 37°C for an hour under a constant shaking (300 rpm) in a 500 ml biotinylation mix with a concentration of 1x ligation buffer, 0.09 mM of dATP-dGTP-dTTP, 0.03 mM of Biotin-14-dCTP and 0.64 U/mL of Klenow fragments. After biotinylation, the samples were incubated for three hours at room temperature in a 1.2 mL ligation reaction with a concentration of 1x ligation buffer, 100 mg/mL of BSA, 1 mM of ATP and 0.4 U/mL of T4 DNA Ligase. The samples were then incubated overnight at 65°C after adding 20μl of 0.5M EDTA, 80μl of 10% SDS and 1.6 mg of Proteinase K. DNA was extracted with 1 volume of phenol/cholorform/isoamyl (24:24:1) alcohol, followed by 30 seconds of vortex at top speed and a five-minute centrifugation at top speed. We precipitated the DNA by adding 1/10 volume of 3M NaAC pH5 and two volumes of cold EtOH 100%, followed by a 30-minute incubation at −80°C and a 20-minute centrifugation at 14,000×g and 4°C. The DNA pellet was washed with 1mL of EtOH 70%, then dried at 37°C for 10 minutes and resuspended in 100μl 1x TE buffer with 1mg/ml of RNase. DNA was sheared to 250-500bp fragments using Covaris S220, purified with AMPure beads (0.6X) (Beckman) and eluted in 20μl 10mM Tris pH8.0. Biotinylated but not ligated DNA fragments were first removed by T4 DNA polymerase treatment (final concentration=300 U/pellet; NEB), and the biotin-labeled fragments were selectively captured by Dynabeads MyOne Streptavidin C1 (Invitrogen). The libraries were prepared using NEB Ultra II library preparation system and sequenced on the NextSeq2000 Illumina platform (2×150 bp) (**Table S21**).

We scaffolded the genomes from Denoeud et al.^26^ into chromosome-level assemblies using the Hi-C data. We filtered the low-quality Hi-C reads using Trimmomatic v0.39^94^ (ILLUMINACLIP:2:30:10 LEADING:25 TRAILING:25 SLIDINGWINDOW:4:15 MINLEN:75 AVGQUAL:28). We mapped the Hi-C reads against each genome assembly using BWA-mem v0.7.17-r1188^95^ as implemented in the Juicer v1.6 pipeline^96^ to generate a contact map, which was then fed to 3D-DNA v190716^97^ to scaffold the genomes into chromosomes. The obtained scaffolds were manually inspected against the contact maps to solve the limits of each chromosome using Juicebox v1.11.08^98^. The PhaeoExplorer gene annotations^26^ were lifted into the new assemblies using Liftoff v1.6.1^99^, while the annotation of transposable elements was performed using RepeatModeler2^100^. We scaffolded the genomes of *Ectocarpus crouaniorum* and *Desmarestia dudresnayi* into chromosomes using a reference-guided assembly with RagTag v2.0.1^101^ against the chromosome-level assemblies of *Ectocarpus* sp. 7 and *Desmarestia herbacea*, respectively.

#### Discovery of the UV sex determination regions

Male sex determining regions (V-SDR) in *S. promiscuus*, *U. pinnatifida*, *D. herbacea* and *D. dichotoma,* as well as female sex determining region (U-SDR) in *D. herbacea* were analyzed following a YGS approach developed by Carvalho and Clark^102^ and coverage analysis described previously^103^. The YGS method principle is to identify male or female sex-linked scaffolds by comparing kmer frequencies between reference genome assembly and kmers generated from DNAseq reads of the opposite sex. Regions in the male reference genome with low density coverage of female kmers will indicate candidate male SDR sequences, similarly, female genomic scaffolds with low coverage in male kmers will denote female SDR region. First, fifteen base pair kmer sequences were generated from respective Illumina reads (**Table S21**) using Jellyfish v2.3.0 count (-m 15 -s 10G -C --quality-start=33 --min-quality=20) and converted to fasta format with Jellyfish dump (--lower-count=5)^104^. Next, non-overlapping 100kb sliding windows of the reference chromosome genome assemblies were created using seqkit v2.3.1^105^ and used as input for the YGS.pl script together with the fasta kmer files produced in the previous step. Genomic windows with a minimum of 70% of unmatched single copy kmers were then retained as candidate male or female SDR sequences. These regions were further validated by the coverage analysis. In detail, the short Illumina reads coming from males and females of each investigated species were trimmed with Trimmomatic^94^ (see above) and mapped to the reference genome, for which the SDR was to be studied, using HISAT2^106^ (default settings). Bam files produced by HISAT2 were used as input for Mosdepth^107^ to calculate coverage in 100kb windows along the genome sequence (-m-n -b 100000 --fast-mode -Q 30). Read mapping depth in genomic windows was normalized by the genome-wide mean for each sex and the coverage in genomic intervals was then compared between males and females. Because V-SDR-linked sequences are present only in males, we expect them to have similar read coverage as autosomal regions in males, but little or no coverage in females (and conversely for the U-SDR sequences). The comparison focused on regions within male reference genomes where the coverage in males fell within the range of 75% to 125% of the genome average, while the coverage in females remained below 50% of the genome average. These findings were then cross-referenced with the results obtained from the YGS analysis. The reverse strategy was applied to female U-SDR regions for a comprehensive evaluation. Both, coverage and kmer analysis, identified identical genomic regions (**Table S1**). Candidate sex-specific scaffolds were further verified by PCR on at least 4 males and 4 females.

#### Genetic mapping and search for the sex chromosome in Fucus serratus

Three different sets of materials were used in this study: (i) twelve male and twelve female field samples hereafter denoted the 24-individual natural population; (ii) 157 sporophyte progeny population derived from a cross between one male sample and one female sample collected from the field and (iii) three male and three female samples collected from the field for whole-genome sequencing. The 157-progeny population and 24-individual natural population were genotyped by double digest RAD sequencing approach (ddRAD-seq). Briefly, individual genomic DNA was digested with the restricted enzymes PstI and HhaI to obtain fragments that were size selected between 400 and 800 bp before sequencing on in Illumina HiSeq 2500 platform (paired-end 2 x 125 bp). See ^108^ for detailed protocol of the ddRAD-seq.

We performed whole-genome sequencing on Illumina HiSeq 2500 (2x 150 bp paired-end) for the three male and three female samples. For ddRAD-seq data, raw reads were cleaned and trimmed with Trimmomatic as above and mapped to the draft genome of *Fucus serratus* male. For the progeny population, genotypes were called from the obtained bam files, using the Stacks pipeline (v2.5)^109^. The obtained vcf files were filtered with VCFtools v0.1.16^110^ and bcftools^111^ (max missing per locus:30%, max missing per sample:40%, max mean coverage:30, minQG:20).

The filtered vcf file of the progeny population was used to construct a genetic map with Lep-MAP3^112^. Briefly, ParentCall2 module was used to call parental genotypes, SeparateChromosomes2 module was used to split the markers into linkage groups and OrderMarkers2 module was used to order the markers within each linkage group using 30 iterations per group and finally computing genetic distances. Phased data were converted to informative genotypes with the script map2genotypes.awk. We used different approaches to identify the SDR in *Fucus serratus*:

##### Coverage analysis

We combined whole-genome sequence data from the three males and three females alongside the ddRAD-seq data of the 24-individual natural population, mapping both datasets to the *F. serratus* male genome assembly using bwa-mem^95^. Coverage analyses have been done in several ways:

– Using SATC (sex assignment through coverage)^113^, a method that uses sequencing depth distribution across scaffolds to jointly identify: (i) male and female individuals, and (ii) sex-linked scaffolds. This identification is achieved by projecting the scaffold depths into a low-dimensional space using principal component analysis and subsequent Gaussian mixture clustering. Male and female whole genome sequences were used for this analysis.
– Using the method SexChrCov described in ^114^ with the 24-individual natural population.
– Using the method DifCover^115^ which identifies regions in a reference genome for which the read coverage of one sample is significantly different from the read coverage of another sample when aligned to a common reference genome. The 24-individual natural population was used for this analysis.
– Using soap.coverage v2.7.9^116^ to calculate the coverage (number of times each site was sequenced divided by the total number of sequenced sites) of each scaffold in each sample. For each scaffold, the male to female (M:F) fold change coverage was calculated as log2(average male coverage) – log2(average female coverage). The 24-individual natural population was used for this analysis.

##### F_ST_ and sex-biased heterozygosity

This approach has been previously used to find sex linked genomic regions in several studies^117,118^. Using the 24-individual natural population, *F*_ST_ was calculated using vcftools^110^. Sexbiased heterozygosity was defined as the log10 of the male heterozygosity:female heterozygosity, where heterozygosity is measured as the fraction of sites that are heterozygous. This ratio is expected to be zero for autosomal scaffolds and elevated on young sex scaffolds due to excess heterozygosity in males.

##### Identification of eventual female scaffolds that failed to map to the male reference genome

Vcftools and bedtools were used to extract female regions that did not map to the reference genome, consistently in the three re-sequenced female samples. All candidate contigs were tested by PCR in 4 males and 4 females.

### Synteny analyses, SDR evolutionary strata and transitions to co-sexuality

Whole-genome synteny comparisons were performed for each pair of chromosome-level assemblies using MCscan v1.2.14^119^, both between different species, between sex chromosomes in the same species and between hermaphrodites and their closest relatives with U/V chromosomes. The putative gametologs between sex chromosomes that were predicted with MCscan were reassessed using OrthoFinder v2.5.4^120^ and best reciprocal DIAMOND v2.1.8.162^121^ hits.

We calculated the number of synonymous substitutions per synonymous site (*Ks*) for each pair of male and female gametologs as a proxy to assess the relative time at which both genes diverged from each other. The amino acid sequences of each pair of gametologs were aligned with MAFFT v7.520^122^ and subsequently aligned into codons using pal2nal v14^123^. The *Ks* values were calculated using the model by Yang & Nielsen^124^ as implemented in KaKs_calculator v2.0^125^.

We evaluated the male or female identity of the genes in the co-sexual species whose orthologs were found within the SDR in their closest non-co-sexual relatives. For this, we compared the results obtained with MCscan^119^ against the orthogroup prediction performed with OrthoFinder^120^, with best reciprocal DIAMOND^121^ hits and by calculating gene trees for each orthogroup using an amino acid alignment with MAFFT^122^ and gene tree reconstructions using FastTree v2.1.11^126^.

### Ancestral reconstruction of the male SDR

We searched for ortholog genes within the V-SDR of five species (*Ectocarpus* sp. 7, *Scytosiphon promiscuus*, *Undaria pinnatifida*, *Desmarestia herbacea* and *Dictyota dichotoma*) in our OrthoFinder results. Once we determined the evolutionary relationship of the genes within the V-SDR, we used the software Count v10.04^127^ to estimate the ancestral content of the V-SDR throughout a phylogeny and determine the most likely scenario of V-SDR evolution in the brown algae. We employed posterior probabilities under a phylogenetic birth-and-death model with independent gain and loss rates across each branch in the phylogeny. We modeled the independent gain and loss rates through 10 gamma categories and performed 1000 optimization rounds with a convergence threshold on the likelihood > 0.1 to find the most fitting model for the data. The branch lengths in the tree that were used for the ancestral state reconstruction were retrieved from the molecular clock analysis performed by ^24^. We distinguished between conserved V-SDR genes that are ancestral and parallel acquisitions of the same gene in the V-SDR by analyzing gene trees between male and female genomes, in addition to female transcriptome assemblies of *Dictyota dichotoma* and *Undaria pinnatifida*. Sequence alignments were done using MAFFT^122^ with default settings and uploaded to http://www.phylogeny.fr/ platform. Alignments were further curated using Gblocks v0.91b^128^ (Min. seq. for flank pos.: 85%, Max. contig. nonconserved pos.: 8, Min. block length: 10). Trees were produced by PhyML v3.1^129^ with default model and visualized in TreeDyn v198.3^130^. Approximate Likelihood-Ratio test (aLRT) was chosen as statistical test for branch support. We inferred the function of the ancestral V-SDR genes through the annotation of genes in *Ectocarpus* sp. 7 belonging to that orthogroup.

### Genomic content across chromosomes

We used closely-related genome assemblies available in the PhaeoExplorer database^26^ to assess the depletion of orthologs in the sex chromosome. We predicted one-to-one orthologs using OrthoFinder^120^ between the following species pairs: *Ectocarpus* sp. 7 with *Ectocarpus siliculosus*, *Scytosiphon promiscuus* with *Chordaria linearis*, *Undaria pinnatifida* with *Saccharina japonica*, *Fucus serratus* with *Fucus distichus*, *Desmarestia herbacea* with *Desmarestia dudresnayi*, and *Dictyota dichotoma* with *Halopteris paniculata* (**Table S22**). We calculated the expected number of detectable orthologs for each chromosome and compared it against the observed number of detected orthologs using chi-squared tests. We performed Benjamini-Hochberg corrections to the *p*-values of the chi-squared tests to control the false discovery rate (FDR) in the analysis^131^.

GenEra^42^ was used by running DIAMOND in ultra-sensitive mode^121^ against the NCBI NR database and all the PhaeoExplorer proteins^26^ to perform a phylostratigraphic analysis (e-value threshold of 10^-5^) and calculate the relative ages of each gene in each genome (**Table S23**). The gene age categories outside of the brown algae and *Schizocladia ischiensis* were based on the taxonomic classification of each species within the NCBI Taxonomy database^132^, while the gene ages within the brown algae were manually assessed to reflect the evolutionary relationships obtained in the PhaeoExplorer maximum likelihood tree^26^. We performed Wilcoxon rank-sum tests in R v4.3.1^133^ to assess nonrandom differences in gene age distributions between pairs of chromosomes (**Table S12**). We performed Benjamini-Hochberg corrections to the *p*-values of the Wilcoxon rank-sum tests to control the FDR in the analysis^131^. The gene ages responsible for these differences were found by evaluating the standardized residuals using mosaic plots (**Figs. S7-S11, S13-S16, S18-S19**). The relative gene ages in Fig. 3B and in **Figs. S7-S11, S13-S16 and S18-S19** were plotted against the chromosome-level assemblies using karyoploteR v1.20.3^134^.

We used the *Ks* values between pairs of species as a proxy for neutral mutation rates across six of the seven chromosome-level assemblies by using the most closely related genome assemblies available in the PhaeoExplorer database^26^. We used the same set of one-to-one orthologs detected between species pairs as for the ortholog-depletion test (**Table S22**). However, the evolutionary distance between *Dictyota dichotoma* and *Halopteris paniculata*prevented us from calculating reliable *Ks* values for this species since synonymous substitutions reached the point of saturation. The amino acid sequences of each pair of orthologs were aligned with MAFFT^122^ and subsequently aligned into codons using pal2nal^123^. The *Ks* values were calculated using the model by Yang & Nielsen^124^ as implemented in KaKs_calculator v2.0^125^. We also evaluated the difference in *Ks* values between the autosomes and the sex chromosomes through FDR-corrected Wilcoxon rank sum tests (**Table S14**). We calculated the protein-coding density, the density of transposable elements and the taxonomic identity of these transposable elements within 100 kb non-overlaping windows across each chromosome using bedtools^91^ (**Table S24**). The differences in protein-coding space, TE content and TE classification between the autosomes and the sex chromosomes were also performed using FDR-corrected Wilcoxon rank sum tests (**Tables S8-S10**). All the genomic features were plotted using shinyCircos-V2.0^135^.

We tested for *de novo* gene birth events in six monoexonic genes contained within the positions 3886599 to 4923391 in chromosome 13 of the *Ectocarpus* sp. 7 genome assembly^27^. We focused on monoexonic genes to facilitate the testing procedure of *de novo* birth events^50^. We searched for protein-coding homologs of these genes by performing a BLASTp^87^ search with an e-value threshold of 10^-3^ against the annotated proteins of ten additional *Ectocarpus* species^26^ (see **Table S21**). We subsequently searched for noncoding regions that were homologous to the candidate genes by performing a tBLASTn^87^ search with an e-value threshold of 10^-3^ against the genome assemblies of the ten *Ectocarpus* species. Once we established the coordinates for the orthologous proteins and noncoding regions in the other *Ectocarpus* species, we extracted the nucleotide sequences from each genome assembly while adding 100 bp upstream and downstream from the BLAST coordinates to encompass possible start and stop codons that might be found in the vicinity of the matched sequences. The nucleotide sequences were initially aligned with MAFFT^122^. We subsequently used the evolutionary relationships between *Ectocarpus* species inferred by Akita et al^136^ to re-align the sequences in a phylogenetically-aware fashion and infer the ancestral state of the sequence at each node in the phylogeny through the maximum likelihood approach implemented in PRANK v170427^137^. We manually assessed the coding potential of the ancestral and extant sequences in the alignment using Aliview v1.27^138^ to infer the enabling mutations that led to the *de novo* birth of these genes from a non-coding background, and whether the non-annotated orthologous sequences found with tBLASTn represent unannotated sequences with coding potential, pseudogenization events or non-coding DNA sequences^50^.

### Gene expression analysis

We used kallisto v.0.44.0^139^ to calculate gene expression levels using 31-base-pair-long k-mers and 1000 bootstraps. Transcript abundances were then summed within genes using the tximport v3.19 package^140^ to obtain the expression level for each gene in TPM. Differential expression analysis was done in DESeq2 v3.19 package^141^ in R v.4.3.1, applying FC>=2 and Padj<0.05 cut-offs. Sex biased gene expression analysis in *Ectocarpus* sp. 7, *Scytosiphon promiscuus*, *Undaria pinnatifida*, *Desmarestia herbacea* and *Dictyota dichotoma* was estimated between mature male and female gametophytes (gametophytes baring reproductive structures). To discover genes with sporophyte biased expression in *Ectocarpus* sp.7, *Scytosiphon promiscuus*, *Undaria pinnatifida* and *Dictyota dichotoma* we first calculated the differential expression between male gametophytes and sporophytes, as well as female gametophytes and sporophytes. Genes that showed significant sporophyte-biased expression (FC>=2, padj<0.05) in both comparisons were considered sporophyte-biased.

A total of 314.2 M RNA-seq reads from *F. vesiculosus* male, female and vegetative tissue were assembled *de novo* with rnaSPAdes^142^ using kmer values of 33 and 49. Assembly quality was assessed by (pseudo)mapping reads back onto the resulting assembly and retaining “good” contigs as defined using TransRate v1.0.3^143^ with default settings. The resulting 159,108 contigs were aligned with BLASTx^87^ against a database of Stramenopile proteins and those with top hits against brown algae (Phaeophyceae) were retained as the final curated reference transcriptome (36,394 contigs, N50 = 1770 bp). Transcript expression levels were determined by mapping the reads from all samples against the reference transcriptome using Bowtie2^144^ and the RSEM-EBSeq v1.3.3^145^ pipeline and relative expression values were recorded as transcripts per million (TPM). All samples used in the gene expression analysis can be found in **Table S21.**

## Notes

### Competing Interest Statement

The authors have declared no competing interest.

### Summary of Updates

We added additional data and analysis (Figure 3B and 3C) and also S12-S17

